# AMPA receptor anchoring at CA1 synapses is determined by an interplay of N-terminal domain and TARP γ8 interactions

**DOI:** 10.1101/2020.07.09.196154

**Authors:** Jake F. Watson, Alexandra Pinggera, Hinze Ho, Ingo H. Greger

## Abstract

AMPA receptor (AMPAR) abundance and positioning at excitatory synapses regulates the strength of transmission. Changes in AMPAR localisation can enact synaptic plasticity, allowing long-term information storage, and is therefore tightly controlled. Multiple mechanisms regulating AMPAR synaptic anchoring have been described, but with limited coherence or comparison between reports, our understanding of this process is unclear. Here, combining synaptic recordings and super-resolution imaging, we compare the contributions of three AMPAR interaction domains controlling transmission at hippocampal CA1 synapses. We show that the AMPAR C-termini play only a modulatory role, whereas the extracellular N-terminal domain (NTD) and PDZ interactions of the auxiliary subunit TARP γ8 are both crucial, and each is sufficient to maintain transmission. Our data support a model in which γ8 accumulates AMPARs at the postsynaptic density, where the NTD further tunes their positioning. This interplay between cytosolic (γ8) and synaptic cleft (NTD) interactions provides versatility to regulate synaptic transmission and plasticity.

## Introduction

Excitatory synaptic transmission is primarily mediated by AMPARs^1^. These glutamate-gated cation channels are concentrated at the postsynaptic density (PSD) to mediate fast neuronal communication. This receptor is also central to synaptic plasticity: activity-dependent changes in the abundance of synaptic AMPARs can bi-directionally modify the strength of transmission, leading to either long-term potentiation (LTP) or long-term depression (LTD). For this reason, the mechanisms controlling the synaptic localisation of AMPARs have been intensively investigated for decades^2,3^.

AMPARs are exocytosed to the cell surface and enter the synapse through lateral diffusion, where they are trapped in the PSD^4–6^. There, they concentrate into sub-synaptic ‘nanoclusters’ of higher density^7,8^, which have been suggested to align with presynaptic vesicle release sites for efficient transmission^9–11^.

AMPAR complexes are assembled from four core subunits and various auxiliary proteins^12^. At hippocampal CA1 synapses, GluA1/2 heteromers predominate and principally associate with the auxiliary subunit TARP γ8 (Transmembrane AMPAR regulatory proteins, **Fig. 1a**)^13–15^. TARPs bind to the major PSD components PSD-93/95, through PDZ (PSD-95, Dlg, ZO-1) interactions of their extreme C-terminus^16–19^. This interaction is currently the best characterized mechanism of AMPAR synaptic localisation.

**Fig. 1.**
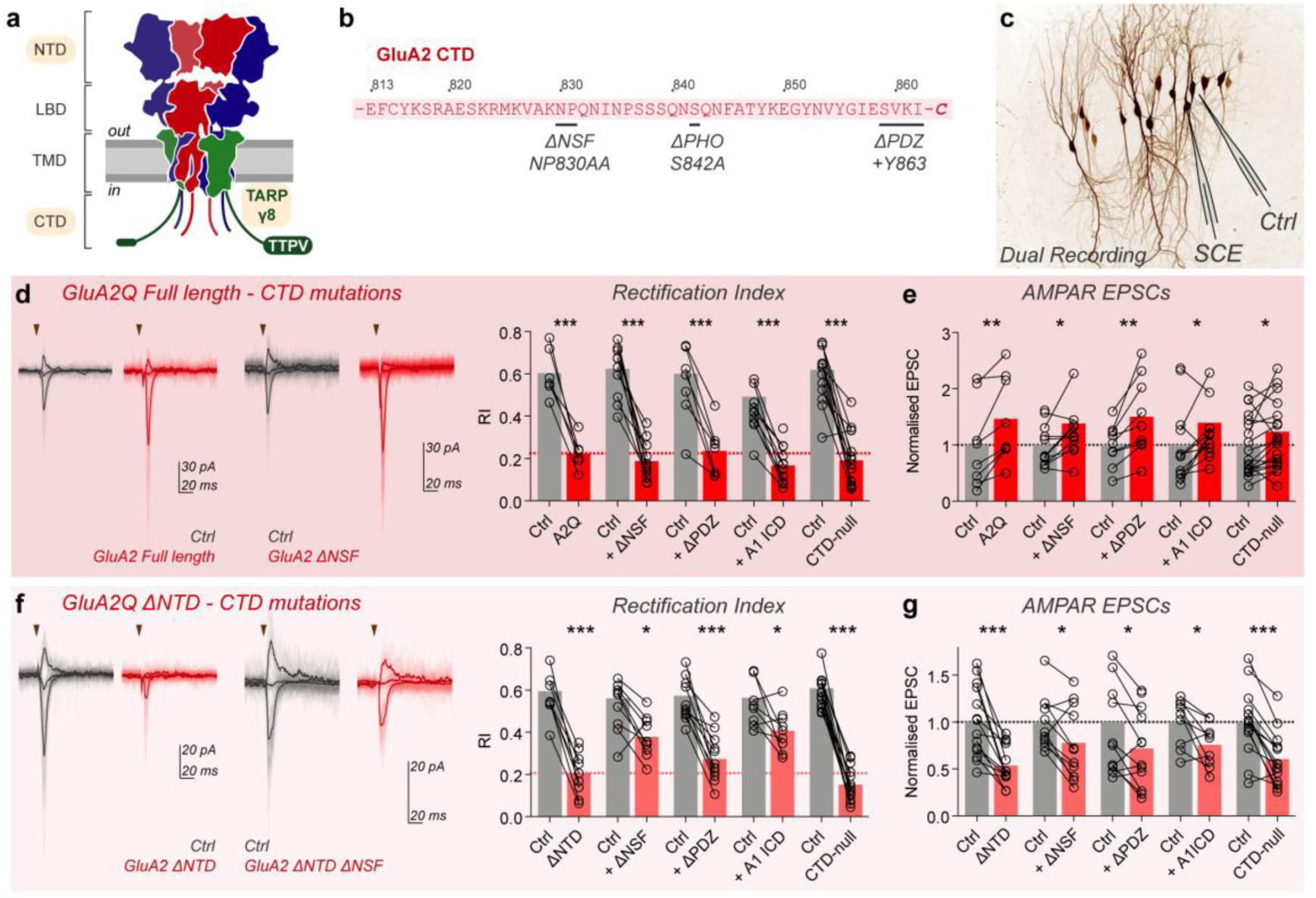
GluA2 CTD interactions exert a modulatory but non-essential influence on receptor synaptic anchoring. **a** Schematic of heteromeric AMPAR architecture (blue - GluA1, red - GluA2) demonstrating the four domain layers (NTD - N-terminal or amino-terminal domain; LBD - ligand binding domain; TMD - transmembrane domain; CTD - C-terminal domain) and association with the auxiliary subunit TARP γ8 (green), with the studied interaction domains highlighted yellow. **b** GluA2 CTD protein sequence with interaction mutations indicated. **c** Dual recording configuration involves simultaneous patch-clamp of untransfected (Ctrl) and single-cell electroporated (SCE) neurons, depicted on fluorescence image of EGFP expressing CA1 neurons after SCE. **d-e** Dual synaptic recordings on GluA2Q construct expression. Full-length GluA2Q robustly reduces synaptic rectification index (**d**) and increases AMPAR EPSC amplitudes (**e**). No CTD modifications altered this phenotype, demonstrating a non-essential role for this domain in synaptic receptor anchoring (**d-e**). Example dual recordings (untransfected cell - Ctrl, grey; transfected cell - red) for GluA2Q with (right) and without (left) ΔNSF mutation are depicted with individual sweeps (light) and average response (bold). Stimulation is indicated as a triangle. GluA2 ΔNTD causes a robust reduction in the rectification index of synaptic responses (**f**) and a reduction in AMPAR EPSC amplitudes (**g**). These changes are partially alleviated by some GluA2 CTD mutations such as NSF site mutagenesis (ΔNSF) or GluA1 intracellular domain (ICD) exchange (**f-g**). Pre-normalised data is presented in **Supplementary Figure 1f-g**, and data values are presented in **Supplementary Table 1**. Bars represent mean values.

The sequence-diverse AMPAR C-terminal domains (CTDs) have multiple phosphorylation and interaction sites for cytosolic proteins (**Fig. 1b, 2a**)^20^, and were central to previous synaptic anchoring models^21,22^. The GluA2 CTD interacts with NSF (N-ethylmaleimide sensitive fusion protein) and the scaffolding proteins GRIP1/ABP and PICK1, with suggested roles in both surface delivery and synaptic anchoring, while the GluA1 CTD interacts with SAP97^21,22^. It was proposed that the GluA1 CTD mediates activity-dependent AMPAR trafficking in LTP, while the GluA2 CTD drives constitutive synaptic delivery^23,24^. This model has since been debated, and the role of AMPAR CTDs remains unclear^25,26^.

Recently, we and others demonstrated a role for the extracellular AMPAR N-terminal domain (NTD; **Fig. 1a**) in synaptic transmission and plasticity^27,28^. This domain, sequence-diverse like the CTD, offers subunit-specific synaptic anchoring through interactions with synaptic cleft proteins and has been implicated previously in receptor clustering by neuronal pentraxins^29,30^.

However, the plethora of proposed interactions and absence of comparison between reports has obscured the core mechanisms of AMPAR synaptic anchoring. Here, we perform a comprehensive comparative analysis of the three major AMPAR interactions, CTD, NTD and TARP γ8 PDZ, to disentangle their roles in CA1 synaptic transmission. We demonstrate that the AMPAR CTDs are not required for synaptic anchoring but play a modulatory role in this process. AMPAR transmission is primarily reliant on intracellular interactions of γ8 with the PSD to accumulate receptors at synaptic sites, and on subunit-specific NTD interactions in the synaptic cleft. An interplay of these two core mechanisms defines the strength of transmission and facilitates the changes that underlie synaptic plasticity.

## Results

### Mutations in the GluA2 CTD do not affect synaptic anchoring

We first set out to clarify the role of the GluA1 and GuA2 CTDs, using single-cell electroporation to express exogenous AMPAR subunits in hippocampal organotypic slices, and dual recordings from transfected and untransfected CA1 pyramidal neurons (**Fig. 1c**; see **Supplementary Figure 1a** for experimental details)^27^. Expression of GluA2, unedited at the Q/R site in the channel pore (GluA2Q), provides a read-out for synaptic localisation, where a change in the rectification index (RI) indicates that exogenous (rectifying homomeric) receptors have replaced endogenous (non-rectifying heteromeric) receptors^23,27^. Exogenous GluA2Q reaches the Schaffer collateral to CA1 synapse, causing a strong reduction in synaptic RI and an increase in AMPAR EPSC amplitudes^27^ (**Fig. 1d-e**; **Supplementary Figure 1f**). We assessed whether CTD interactions were necessary for this anchoring by mutating known CTD interaction sites on the GluA2Q construct (**Fig. 1b; Supplementary Figure 1b**)^21^. These include mutating the NSF interaction site (NP830AA^24^), blocking PDZ-interactions with ABP/GRIP or PICK1 (addition of Y863^24^), exchanging the intracellular domains (ICD - CTD and loop 1) of GluA2 with that of GluA1 to prevent any interactions that could mediate constitutive GluA2 delivery^24^, and simultaneously mutating major protein interaction and phosphorylation sites in the GluA2 CTD^21^ (CTD-null). All constructs trafficked to the cell surface, as measured by RI changes in somatic outside-out patches (**Supplementary Figure 1c-d**). However, none of these mutations altered or prevented synaptic expression of GluA2Q. All constructs reduced the synaptic RI (**Fig. 1d**) and increased EPSC amplitudes similarly to full-length GluA2Q (**Fig. 1e**), suggesting that GluA2 CTD interactions are not essential for AMPAR synaptic delivery.

### CTD interactions modulate synaptic localisation of GluA2 ΔNTD

We previously reported that removal of the GluA2 NTD (GluA2 ΔNTD) limited AMPAR anchoring at the synapse^27^. GluA2Q ΔNTD receptors (unedited at the Q/R site) were present at synapses, as measured by a change in the synaptic RI, however, the amplitude of evoked EPSCs was decreased by ∼50 % relative to untransfected cells (**Fig. 1f-g**; **Supplementary Figure 1g**). This phenotype suggests that while AMPAR anchoring is impaired by NTD removal, GluA2Q ΔNTD also prevents endogenous receptors from maintaining full transmission, likely by competing for limiting interactors.

As the exogenous receptors showed no change in auxiliary subunit association in comparison to endogenous receptors^27^, we assessed whether CTD-interactions cause this effect. Loss of critical protein interactions should alleviate competition with endogenous AMPARs, rescuing both rectification and EPSC depression. Surface trafficking of all GluA2 ΔNTD CTD mutants was unimpaired (**Supplementary Figure 1e**). Neither mutation of the NSF interaction site, nor blocking PDZ-interactions completely prevented GluA2Q ΔNTD detection at synapses, yet NSF-mutation partially alleviated the magnitude of both RI and EPSC amplitude changes (**Fig. 1f-g**), implicating a role for this interaction in synaptic delivery. Replacing the ICD of GluA2Q ΔNTD with that of GluA1 also reduced the effect of GluA2Q ΔNTD on synaptic transmission (**Fig. 1f-g**), further supporting a modulatory role for the GluA2 CTD in receptor synaptic delivery. However, GluA2Q ΔNTD CTD-null caused a decrease in both synaptic RI and EPSC amplitudes, equivalent to GluA2 ΔNTD (**Fig. 1f-g**), indicating that we are yet to fully understand the ways by which GluA2 CTD interacting proteins bind or exert their function.

### GluA1 CTD interactions are not essential for synaptic anchoring

We extended this analysis to GluA1, which is associated with activity-dependent AMPAR trafficking^23,31–33^. When expressed alone, GluA1 constitutively trafficked to synaptic sites, as evidenced by a change in the synaptic RI (**Fig. 2b**, see also **Supplementary Figure 2b**)^27,28^. Previous work highlighted an essential role for the GluA1 PDZ ligand in synaptic potentiation^23,24^, therefore we mutated this site in GluA1 (T887A) (**Fig. 2a**). PDZ mutation did not prevent trafficking to the cell surface (**Supplementary Figure 2a**) or synapse (**Fig. 2b**), and co-mutation of the PDZ-ligand and two CTD phosphorylation sites (**Fig. 2a**)^34^, also failed to inhibit GluA1 synaptic localisation (**Supplementary Figure 2c**). Thus, like for GluA2 full-length receptors, we observe no essential role for the GluA1 CTD in constitutive synaptic localisation. Constitutive trafficking is also not dependent on slice activity, as chronic treatment with the Na-channel blocker tetrodotoxin (TTX) did not prevent GluA1-dependent changes to synaptic RI. However, N-terminally tagging GluA1 with GFP prevented it from contributing to transmission (**Supplementary Figure 2d**), in line with previous observations ^24,25^, possibly due to masking of NTD-interaction sites.

**Fig. 2.**
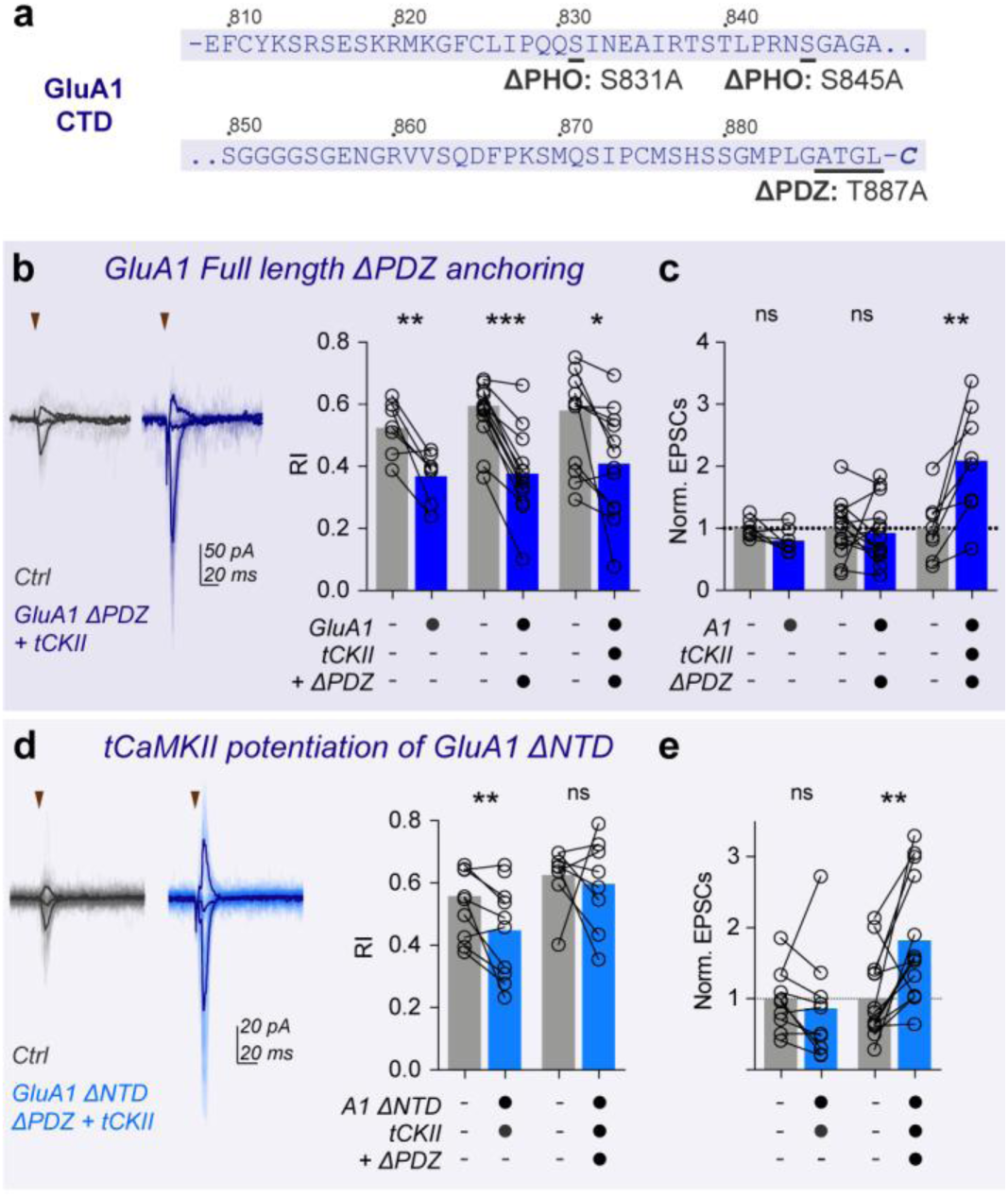
GluA1 CTD interactions exert a modulatory but non-essential influence on receptor synaptic anchoring. **a** GluA1 CTD protein sequences with interaction mutations indicated. Dual recording of RI (**b**) and AMPAR EPSCs (**c**) demonstrates that PDZ-ligand mutation does not prevent full-length GluA1 synaptic anchoring in basal conditions, or in tCaMKII (tCKII)-dependent potentiation. **d-e** tCaMKII (tCKII) potentiation of synaptic responses is prevented by GluA1 ΔNTD, but not by GluA1 ΔNTD with a mutated PDZ-ligand (**d** - Dual cell Rectification Index, **e** - Normalised AMPAR EPSC amplitudes). Pre-normalised data is depicted in **Supplementary Figure 2b**, and data values are presented in **Supplementary Table 2**. Bars represent mean values.

We next assessed the PDZ-dependence of GluA1 in synaptic potentiation, by co-expressing constitutively active CaMKII (truncated (t)-CaMKII)^23^. Similarly to full-length GluA1-expressing cells^23,27^, we observed robust tCaMKII-induced potentiation of GluA1 ΔPDZ-expressing cells, and a significant reduction in RI (**Figure 2b-c**), demonstrating unimpaired incorporation of GluA1 ΔPDZ at synaptic sites upon plasticity. We therefore see no essential requirement for the PDZ-ligand of GluA1 in synaptic transmission or potentiation.

### CTD interactions influence GluA1 ΔNTD in synaptic plasticity

Contrary to GluA2, NTD removal prevents constitutive anchoring of GluA1 and also inhibits maintenance of LTP^27,28^. Moreover, while co-expression of GluA1 full-length with tCaMKII potentiated synaptic EPSCs, expression of GluA1 ΔNTD blocked this potentiation (**Fig. 2e; Supplementary Figure 2b**)^27^. This observation is reminiscent of GluA2 ΔNTD’s dominant negative effect on basal synaptic responses, and it is therefore possible that GluA1 ΔNTD is preventing endogenous receptors from effecting CaMKII potentiation, by competing for GluA1 CTD interactors.

Indeed, while GluA1 ΔNTD-expressing cells exhibited no synaptic potentiation on tCaMKII co-expression, PDZ mutation alleviated this inhibition, and robust tCaMKII-mediated potentiation of current amplitude was observed (**Fig. 2d-e**). While both receptors reached the cell surface (**Supplementary Figure 2a**), neither GluA1 ΔNTD or GluA1 ΔNTD ΔPDZ changed synaptic rectification under basal conditions (**Supplementary Figure 2e**). Together, this data supports some role for the GluA1 PDZ ligand in synaptic potentiation. In summary, our comparative analysis of NTD and CTD interactions demonstrates a far more influential requirement for NTD interactions in AMPAR synaptic localisation, and a non-essential, but modulatory role for the CTD.

### γ8 PDZ interactions accumulate AMPARs at the PSD

The TARP γ8 PDZ interaction with PSD-95 is currently the principal model for AMPAR incorporation into CA1 synapses. We sought to compare intracellular TARP interactions and extracellular NTD interactions in AMPAR synaptic localisation. Using neonatal injection of AAV expressing Cre-EGFP in conditional AMPAR knockout mice (Gria1-3^fl/fl, 13^), we produced organotypic slices with a mosaic of unmodified and AMPAR-null CA1 neurons (**Fig. 3a**). 14 days after Cre transduction, EGFP+ cells showed virtually no excitatory synaptic transmission at -60 mV holding potential, demonstrating successful knockout of all AMPAR subunits expressed in these neurons (**Fig. 3b**). The remaining ∼5-10 % of current was NMDAR-meditated^28^, and NMDAR currents were unaffected by AMPAR deletion when measured at +40 mV holding potential (**Supplementary Figure 3a)**. On this AMPAR-null background, we restored specific AMPAR complexes by single cell-electroporation.

**Fig. 3.**
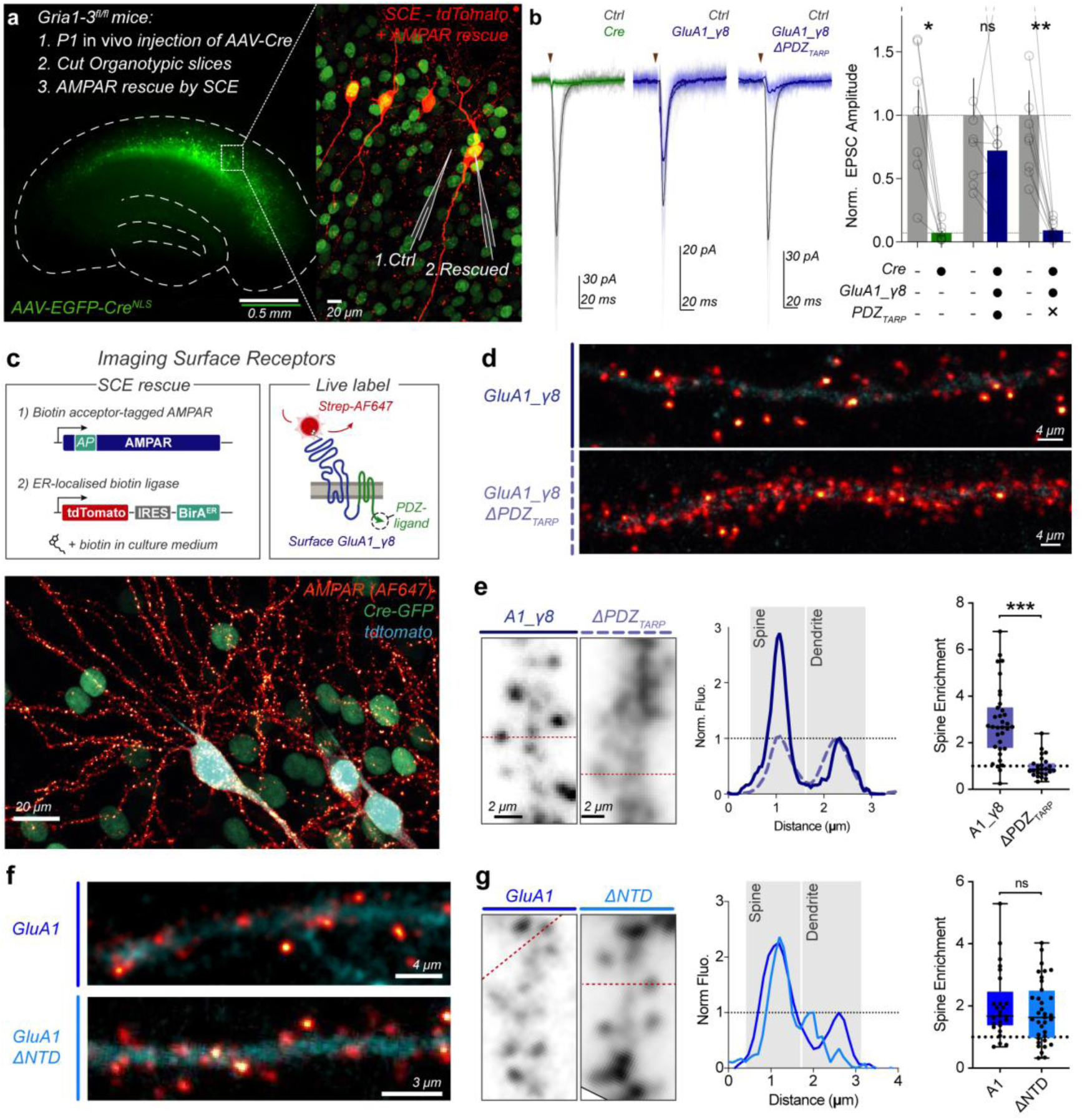
TARP γ8 PDZ-ligand interactions accumulate AMPARs at synaptic sites. **a** AMPAR knockout and rescue strategy in Gria1-3^fl/fl^ tissue. AMPARs in *in vivo* AAV-EGFP-Cre transduced organotypic hippocampal slices (left, outline depicted dashed) are rescued by SCE (right) with coexpressed tdTomato for cell identification. Dual recordings are performed between unmodified (Ctrl) and rescued (EGFP & tdTomato positive) neurons. **b** Cre-transduction abolishes AMPAR synaptic transmission (*Ctrl:* 1.00 ± 0.20, *Cre:* 0.07 ± 0.03, n=7 pairs, p=0.016). GluA1_γ8 transfection rescues synaptic transmission (*Ctrl:* 1.00 ± 0.29, *GluA1_γ8 rescue*: 0.72 ± 0.20, n = 8 pairs, p=0.25), however γ8 PDZ mutation prevents this rescue (*Ctrl:* 1.00 ± 0.20, *GluA1_γ8 ΔPDZ rescue:* 0.09 ± 0.02, n = 9 pairs, p=0.0039). Example traces are depicted with stimulation indicated (triangle). **c** Strategy for fluorescent labelling of surface AMPARs on rescue of knockout neurons using streptavidin labelling of biotinylated AMPARs (left), including schematic of AMPAR-TARP tandem configuration (right). ER-localised BirA biotinylates the acceptor peptide (AP) on the receptor N-terminus. Example image (bottom) of surface AMPARs (red) in Cre-transduced (green) slices after co-rescue with tdTomato cell marker (blue). **d** GluA1_γ8 receptors accumulate in spines along the dendrite, while TARP PDZ mutation causes redistribution of receptors to the dendritic surface. **e** Line profiles (left) of AMPAR fluorescence across spine and dendrite demonstrate enrichment of PDZ-containing receptors in spines. Quantification of receptor distribution (right) shows accumulation of GluA1_γ8 but not GluA1_γ8 ΔPDZ in spines (*GluA1_γ8:* 2.86 ± 0.26 (mean ± SEM), n = 34 spines over 7 cells; *GluA1_γ8 ΔPDZ:* 0.97 ± 0.09, n = 25 spines over 5 cells; p<0.0001). **f** NTD-deletion of GluA1 (GluA1 ΔNTD) did not prevent AMPAR accumulation in dendritic spines as seen by surface receptor imaging. **g** Line profiles (left), and spine enrichment quantification (right) demonstrate spin localisation of GluA1 ΔNTD (*GluA1:* 2.02 ± 0.22 (mean ± SEM), n = 25 spines over 5 cells; *GluA1 ΔNTD:* 1.80 ± 0.17, n = 35 spines over 7 cells; p<0.52).

The ‘TARP-tandem’ configuration, where the AMPAR C-terminus is conjugated to the γ8 N-terminus by in frame expression (termed GluAX_γ8)^35^, allows control over γ8 PDZ-ligand interactions. Expressing GluA1_γ8 in AMPAR-null cells rescued AMPAR synaptic currents, however removal of the C-terminal PDZ-ligand of γ8 (TTPV motif) completely prevented this rescue (**Fig. 3b; Supplementary Figure 3b-c**), in agreement with Sheng et al.^36^.

We and others previously demonstrated that NTD-removal causes a similar attenuation of EPSC rescue in Gria1-3^fl/fl^ neurons^27,28^, therefore both TARP and NTD interactions play major roles in synaptic AMPAR accumulation, even in the absence of endogenous AMPARs. Given that both components are important for functional transmission, we investigated whether they have redundant or parallel roles.

We imaged the distribution of surface GluA1_γ8 or GluA1_γ8 ΔPDZ receptors, containing a N-terminal biotin acceptor peptide sequence (AP), in AMPAR-null cells of organotypic slices upon live-labelling with Streptavidin-AF647^5^ (see procedure in **Fig. 3c**). GluA1_γ8 was distributed throughout the dendritic arbour of CA1 pyramidal cells, with a strong accumulation in dendritic spines (**Fig. 3d**-**e**; **Supplementary Figure 3d-e**). GluA1_γ8 ΔPDZ, showed a dramatic re-distribution, with receptors no longer enriched in spines (**Fig. 3d**-**e**; **Supplementary Figure 3d-e**). The 3-fold enrichment of GluA1_γ8 in spines relative to the dendritic shaft was lost on deletion of the γ8 PDZ-ligand (**Fig. 3e**), explaining the dramatic reduction of transmission (**Fig. 3b**). Therefore, TARP - PSD-95 interactions accumulate receptors in spines, most likely at the PSD.

We next compared the distribution of surface GluA1 and GluA1 ΔNTD. Unexpectedly, both constructs showed a punctate, spine-enriched distribution, suggesting that gross spine accumulation is not dramatically impaired by NTD deletion (**Fig. 3f-g;** see also **Fig. 4d**). Therefore, the functional effect of NTD deletion may occur on a subsynaptic scale.

**Fig. 4.**
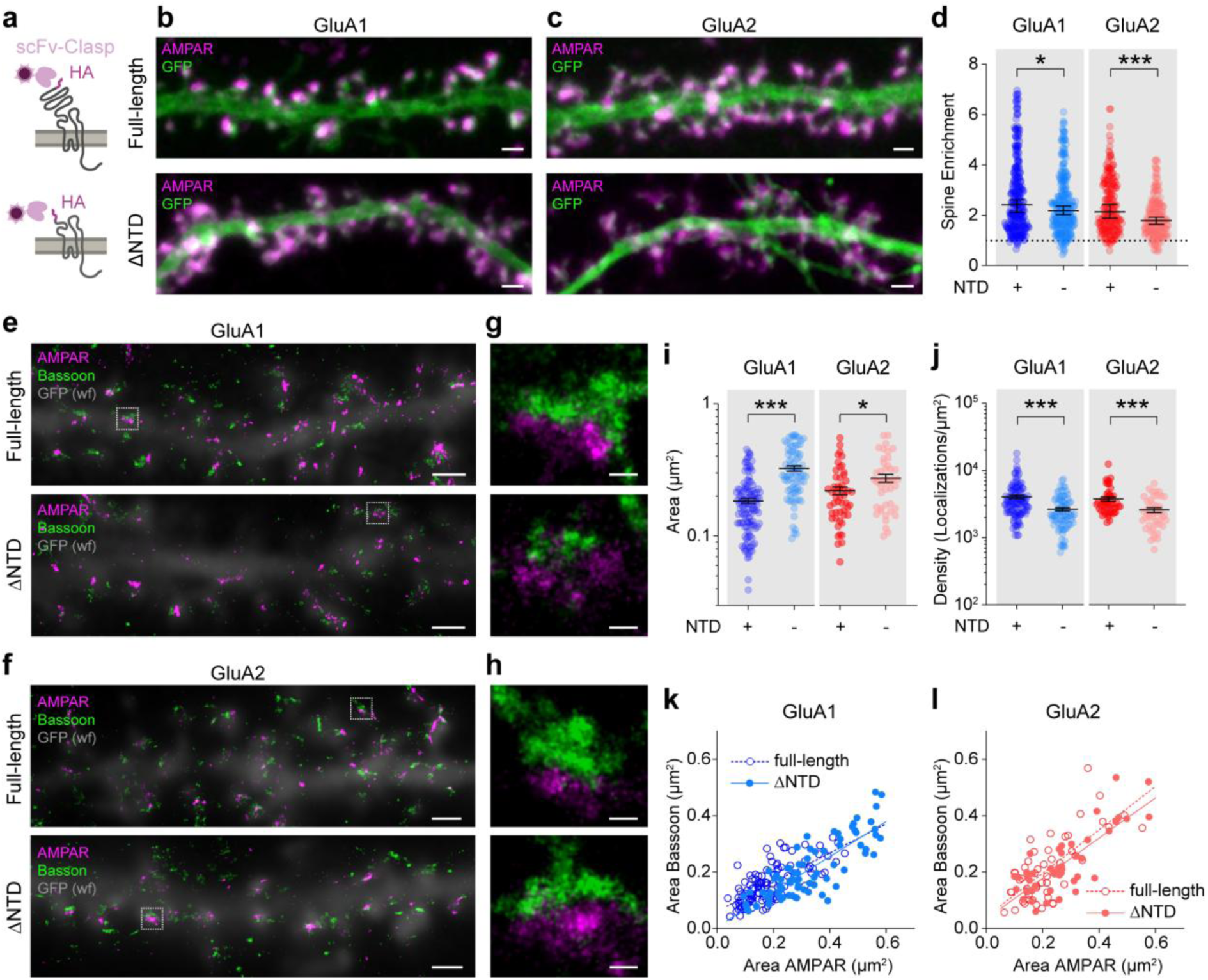
Synaptic distribution of NTD-deleted receptors. **a** Exogenous surface HA-tagged receptors in cultured hippocampal neurons were live-labelled with fluorescently conjugated single chain Fv-Clasp (ScFv-Clasp) avoiding artificial protein cross-linking. **b-c** Representative confocal images of full-length (top) and ΔNTD (bottom) GluA1 (**b**) and GluA2 (**c**) (AMPAR – magenta, GFP – green, Scale bars: 1 µm). **d** ΔNTD receptors were slightly de-enriched at dendritic spines (spine/dendrite fluorescence, left, *GluA1 full-length*: 2.42 ± 2.11-2.63, n=294 spines/16 cells; *GluA1 ΔNTD*: 2.18 ± 2.03-2.37, n=267 spines/16 cells, 5 preparations; p=0.0473, Mann-Whitney test; right, *GluA2 full-length*: 2.14 ± 1.90-2.44, n=200 spines/14 cells; *GluA2 ΔNTD*: 1.79 ± 1.65-1.93, n=187 spines/12 cells, 4 preparations; p<0.0001, Mann-Whitney test). **e-h** Representative 3D STORM images of full-length (top) and ΔNTD (bottom) GluA1 (**e**) and GluA2 (**f**) receptors (magenta) co-labelled with presynaptic marker bassoon (green). Widefield (wf) GFP images overlayed in grey (Scale bars: 1 µm). **g-h** Magnified synaptic clusters within boxed areas in e-f (Scale bars: 100 nm). **i** ΔNTD GluA1 (areas (µm^2^), left, *GluA1 full-length:* 0.183 ± 0.009, *GluA1 ΔNTD:* 0.325 ± 0.015; p<0.0001, unpaired t-test) and ΔNTD GluA2 receptors (areas (µm^2^), right, *GluA2 full-length: 0*.*220* ± 0.015, *GluA2 ΔNTD:* 0.274 ± 0.019; p<0.0193, unpaired t-test) occupied larger synaptic areas than respective full-length controls upon 3D STORM imaging. **j** NTD deletion in both, GluA1 (densities (Localisations/µm^2^), left, *GluA1 full-length*: *4087* ± 248.8, *GluA1 ΔNTD:* 2648 ± 164.6; p<0.0001, unpaired t-test) and GluA2 (densities (Localisations/µm^2^), right, *GluA2 full-length*: *3787* ±2 76.4, *GluA2 ΔNTD:* 2590 ± 208.5; p<0.0001, unpaired t-test) reduced receptor densities at the synapse. **k-l** Post-synaptic AMPAR areas correlated with pre-synaptic bassoon areas (**k**, slope ± S.E. *GluA1 full-length*: 0.5175 ± 0.049 (R^2^ 0.5024), *GluA1 ΔNTD*: 0.6235 ± 0.049 (R^2^ = 0.6768), p=0.1268, F-test: F (DFn, DFd) = 2.353 (1, 189)) and GluA2 ΔNTD (**l**, slope ± S.E. *GluA2 full-length*: 0.7698 ± 0.1056 (R^2^ 0.5202), *GluA2 ΔNTD*: 0.7155 ± 0.082 (R^2^ = 0.6435), p = 0.6857, F-test: F (DFn, DFd) = 0.1648 (1, 91)). Data shown as median ± lower-upper 95% CI (panel d) or mean ± S.E.M (panels i-j). For n-numbers for STORM data see legend to Figure 5.

### NTD-deletion alters the nanoscale arrangement of synaptic AMPARs

To investigate the effect of NTD-deletion on receptor localisation in greater detail, we set up a receptor labelling approach in dissociated hippocampal cultures that allows super-resolution imaging of surface AMPARs. GluA1 and GluA2Q both with and without their NTD were tagged at their N-termini with an HA-epitope, which can be labelled live using a fluorescently-labelled anti-HA single-chain Fv-fragment (scFv-Clasp) **(Fig. 4a**)^37^. This small, monovalent probe (∼5 nm diameter) prevents receptor clustering by epitope cross-linking and increases localisation precision in single-molecule imaging.

Using confocal microscopy, we observed a moderate decrease in surface expression of GluA2 ΔNTD receptors, but not of GluA1 ΔNTD, while intracellular expression was increased for both constructs (**Fig. 4b, c; Supplementary Figure 4a-b**). As seen in slices, the surface-expressed ΔNTD receptors were enriched at dendritic spines, albeit displaying a slightly decreased spine enrichment in comparison to full-length receptors (**Fig. 4d**; **Supplementary Figure 4c-d**).

Since synaptic AMPARs concentrate in ‘nanoclusters’^7,8^, we used 3D STORM (Stochastic Optical Reconstruction Microscopy) to investigate the impact of the NTD on synaptic and sub-synaptic receptor distribution at the nanoscale (**Fig. 4e-l, Fig. 5**). For this experiment, we only analysed neurons displaying comparable AMPAR surface expression levels, as quantified from widefield images collected prior to STORM imaging. Interestingly, both GluA1 and GluA2 ΔNTD mutants occupied a larger synaptic area compared to the respective full-length receptors, which coincided with an increase in the corresponding presynaptic area, marked by the active zone protein bassoon (**Fig. 4i-l**). As the number of observed localisations was only marginally increased for GluA1 ΔNTD but not for GluA2 ΔNTD, this resulted in a significant reduction in the overall density of ΔNTD receptors at the synapse (**Fig 4j and Supplementary Figure 4e**).

**Fig. 5.**
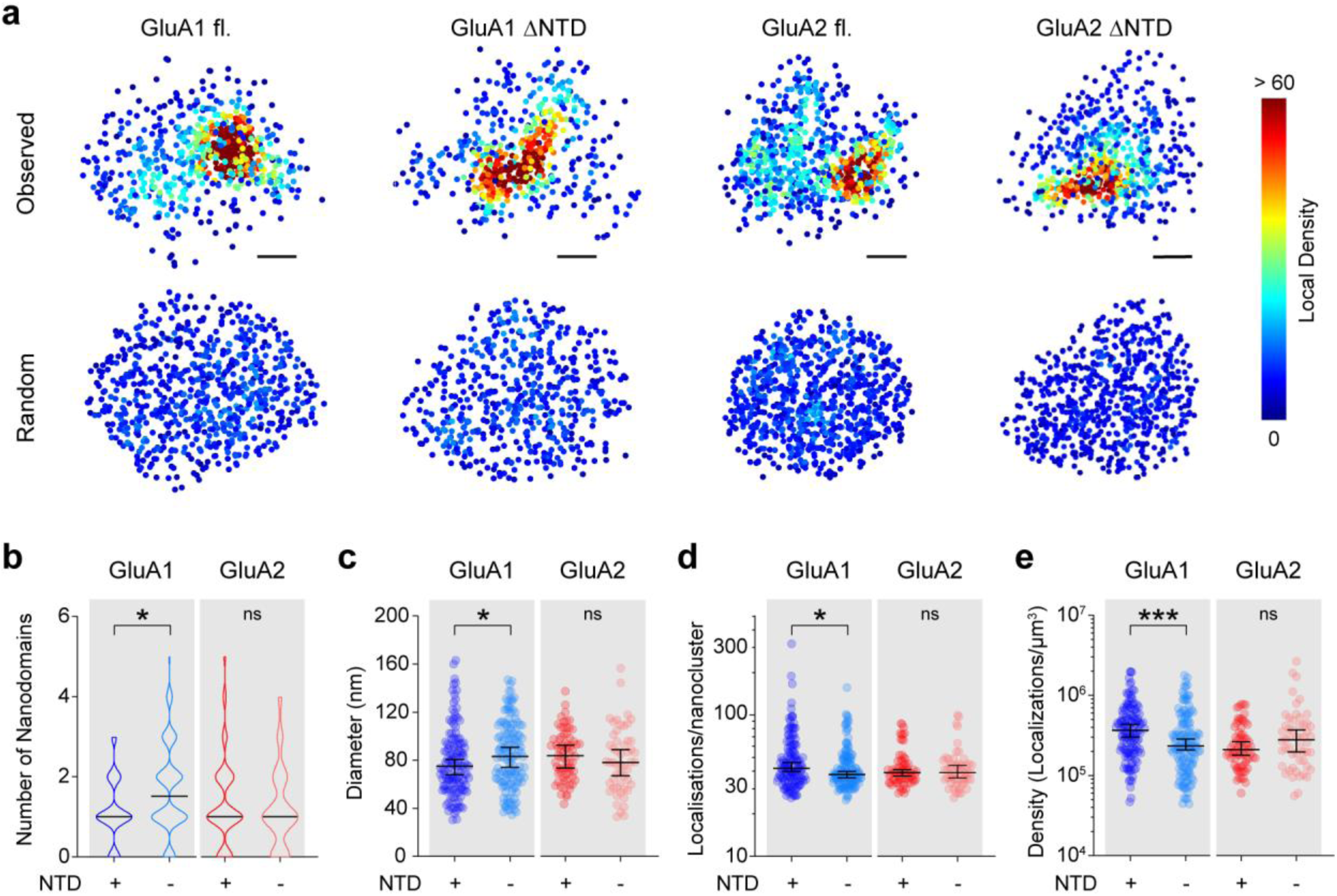
Visualising the sub-synaptic distribution of NTD-deleted receptors using 3D STORM imaging. **a** Representative density plots of synaptic AMPAR clusters reconstructed as scatter plots, highlighting the non-uniform distribution of receptors (cold *vs*. hot colours corresponding to low *vs*. high densities). Observed organisation (top), versus random (bottom) re-distributions of measured localisations shown for comparison (Scale bars: 100 nm). **b** GluA1 ΔNTD receptors exhibited a small increase in nanocluster number per synapse compared to GluA1 full-length (numbers as median ± C.I., left, *GluA1 full-length:* 1 ± 1-1, *GluA1 ΔNTD:* 1.5 ± 1-2, p=0.018, Mann-Whitney test), whereas GluA2 nanocluster number was unaffected by NTD deletion (right, *GluA2 full-length:* 1 ± 1-1, *GluA2 ΔNTD:* 1 ± 1-1, p=0.4895, Mann-Whitney test). **c** Nanocluster diameters (longest axis) were increased for GluA1 ΔNTD (diameters (nm), left *GluA1 full-length:* 75.1 ± 68.0-80.6, *GluA1 ΔNTD:* 83.3 ± 74.1-90.8, p=0.0459, Mann-Whitney test) but unchanged for GluA2 ΔNTD receptors (diameters (nm), right *GluA2 full-length:* 83.7± 73.7-92.6, *GluA2 ΔNTD:* 78.1 ± 67.2-88.9, p=0.2378, Mann-Whitney test) relative to respective full-length controls. **d** GluA1 ΔNTD nanoclusters consisted of fewer localisations compared to GluA1 full-length receptors (left, *GluA1 full-length:* 42.0 ± 40.0-46.0, *GluA1 ΔNTD:* 38 ± 36.0-40.0, p=0.0365, Mann-Whitney test). The number of localisations per nanocluster was unchanged for GluA2 full-length vs ΔNTD (right, *GluA2 full-length:* 39.0 ± 37-41, *GluA2 ΔNTD:* 39.0 ± 36-44, p=0.9189, Mann-Whitney test). **e** NTD deletion of GluA1 (density (localisations/nanocluster), left, *GluA1 full-length:* 366166 ± 301447-436647, *GluA1 ΔNTD:* 235946 ± 209787-286802, p=0.0005, Mann-Whitney test) but not GluA2 (density (localisations/nanocluster), right *GluA2 full-length:* 211710 ± 179581-264751, *GluA2 ΔNTD:* 280631 ± 196833-370233, p=0.1766, Mann-Whitney test) reduced receptor densities within nanoclusters. Data shown as median ± lower-upper 95% CI. STORM data were obtained from 3 culture preparations for GluA1 (*GluA1 full-length:* 113 synapses/12 cells, *GluA1 ΔNTD:* 80 synapses/16 cells) and from 4 preparations for GluA2 (*GluA2 full-length:* 51 synapses/9 cells, *GluA2 ΔNTD*: 44 synapses/10 cells).

To quantify subsynaptic receptor distributions (‘*AMPAR Nanoclusters*’) we used a clustering algorithm (OPTICS) which performs well for datasets with variable densities^38–40^. AMPARs lacking the NTD still assembled into nanoclusters, but exhibited subunit-selective differences (**Fig. 5a**). Specifically, GluA1 ΔNTD nanoclusters displayed a larger diameter, fewer localisations and a reduced cluster density compared to full-length GluA1, while the properties of GluA2 ΔNTD nanoclusters appeared surprisingly unchanged (**Fig. 5c-e**). In addition, we computed that, on average, one nanocluster is present per synapse for GluA1 and GluA2 full-length as well as for GluA2 ΔNTD receptors whereas the averaged cluster number for GluA1 ΔNTD was slightly increased to 1.5 (**Fig. 5b and Supplementary Figure 4f-g**); representative subsynaptic AMPAR distributions are shown in **Fig. 4g, h and Fig. 5a**). Together, these results indicate that the NTD is not essential for nanocluster formation but affects synaptic receptor densities and nanocluster properties in a subunit-specific manner.

### The GluA2 NTD is a minimal unit for synaptic transmission

Subunit-specific effects of the NTD were apparent from our previous work and domain-swap experiments further demonstrated that the GluA2 NTD imparts greater synaptic incorporation than that of GluA1^27^. To shed light on this behaviour, we extended our comparison of TARP and NTD interactions to GluA2. GluA2Q_γ8 robustly rescues evoked AMPAR EPSCs in AMPAR-null cells (**Fig. 6a, 6g**). However, in stark contrast to GluA1_γ8 lacking the γ8 PDZ ligand (**Fig. 3b**), we observed synaptic rescue by GluA2_γ8 ΔPDZ, maintaining around half the relative current amplitude of GluA2Q_γ8 (**Fig. 6b, 6g**). NMDAR currents were unchanged by these manipulations (**Supplementary Figure 5a**).

**Fig. 6.**
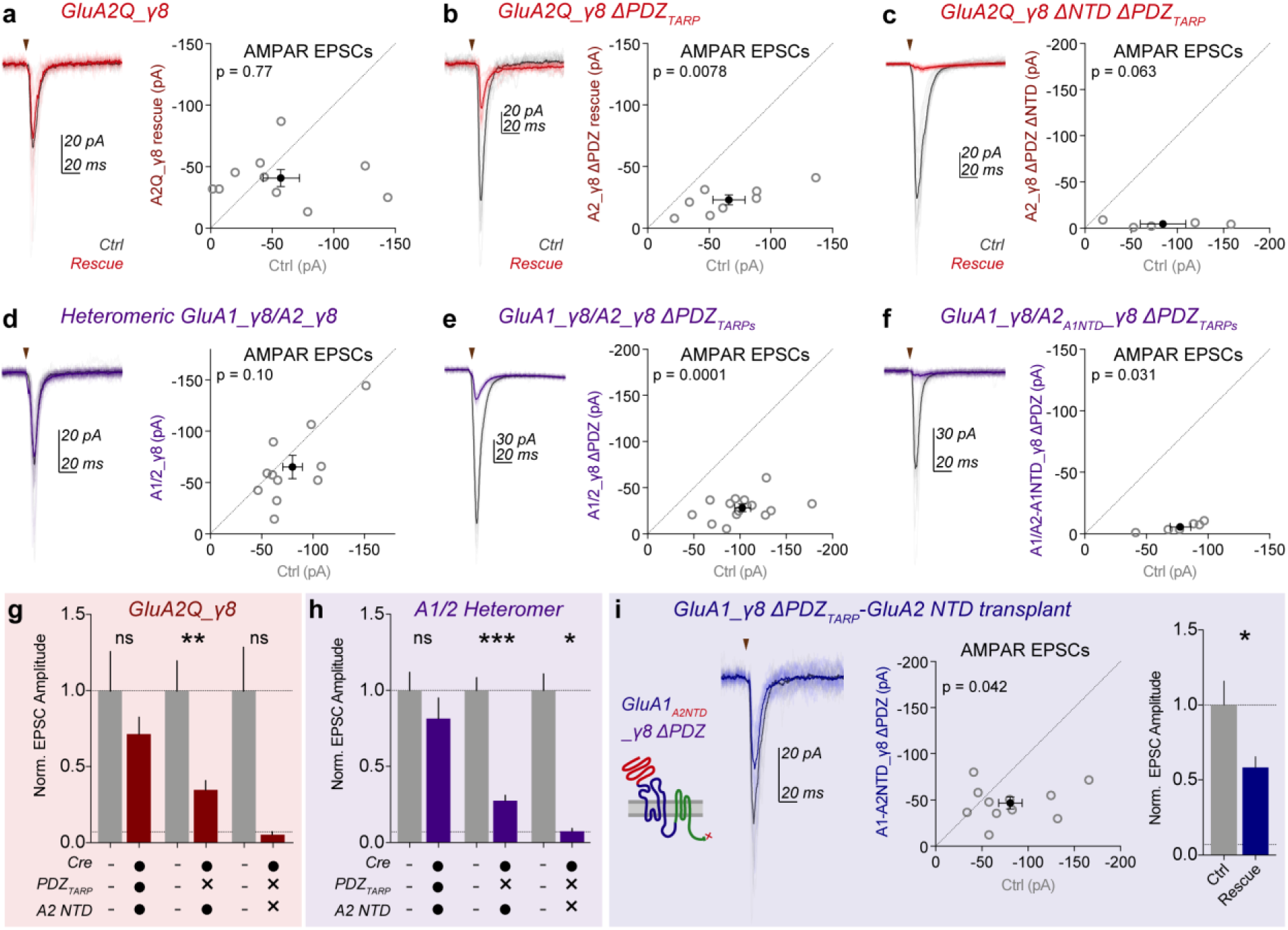
Both γ8 PDZ and GluA2 NTD interactions are sufficient for AMPAR synaptic transmission. **a** Rescue of AMPAR-null cells by GluA2Q_γ8 produces robust synaptic transmission (*Ctrl:* -57.2 ± 15.0 pA, *GluA2Q_γ8 rescue:* -40.9 ± 6.4 pA, n = 10 pairs, p=0.77). **b** γ8 PDZ deletion reduces, but does not prevent AMPAR transmission (*Ctrl:* -65.9 ± 13.1 pA, *GluA2Q_γ8 ΔPDZ rescue:* -22.9 ± 4.0 pA, n = 8 pairs, p=0.0078). **c** Removal of both the γ8 PDZ ligand and GluA2 NTD prevents rescue of AMPAR synaptic transmission (*Ctrl:* -84.4 ± 24.5 pA, *GluA2Q_γ8 ΔPDZ ΔNTD rescue:* -4.5 ± 1.4 pA, n = 5 pairs, p=0.63). **d** Robust transmission is observed on rescue of AMPAR knockout by heteromeric GluA1_γ8/GluA2_γ8 receptors (*Ctrl:* -80.0 ± 9.6 pA, *A1_γ8/A2_γ8 rescue:* -65.2 ± 10.9 pA, n = 11 pairs, p=0.10). **e** γ8 PDZ deletion from both GluA1 and GluA2 rescue constructs reduces but does not prevent AMPAR transmission (*Ctrl:* -102.7 ± 8.7 pA, *A1_γ8/A2_γ8 ΔPDZs rescue:* -28.4 ± 3.6 pA, n = 14 pairs, p=0.0001). **f** Exchanging the GluA2 NTD for that of GluA1 to remove GluA2 NTD from heteromeric AMPAR rescue prevents any rescue of transmission (*Ctrl:* -77.4 ± 8.5 pA, *A1_γ8/A2*_*A1NTD*_*_γ8 ΔPDZs rescue:* -5.8 ± 1.5 pA, n = 6 pairs, p=0.031). Normalised GluA2Q_γ8 (**g**) and heteromeric (**h**) rescue experiments (**g-** *Ctrl:* 1.00 ± 0.26, *GluA2Q_γ8 rescue:* 0.72 ± 0.11; *Ctrl:* 1.00 ± 0.20, *GluA2Q_γ8 ΔPDZ rescue:* 0.35 ± 0.06; *Ctrl:* 1.00 ± 0.29, *GluA2Q_γ8 ΔPDZ ΔNTD rescue:* 0.05 ± 0.02. **h -** *Ctrl:* 1.00 ± 0.12, *A1_γ8/A2_γ8 rescue:* 0.81 ± 0.14; *Ctrl:* 1.00 ± 0.08, *A1_γ8/A2_γ8 ΔPDZs rescue:* 0.28 ± 0.04; *Ctrl:* 1.00 ± 0.11, *A1_γ8/A2*_*A1NTD*_*_γ8 ΔPDZs rescue:* 0.07 ± 0.02). Transmission levels in Cre only expressing cells are depicted as a line. **i** The GluA2 NTD drives synaptic anchoring of GluA1_γ8 ΔPDZ receptors (Amplitudes *-Ctrl:* -80.7 ± 12.8 pA, *GluA1*_*A2NTD*_*_γ8 ΔPDZ:* -47.1 ± 5.8 pA, n = 11 pairs, p=0.042. Normalised data *-Ctrl:* 1.00 ± 0.16, *GluA1*_*A2NTD*_*_γ8 ΔPDZ:* 0.58 ± 0.07).

This result is surprising, given the complete dependence of GluA1_γ8 on γ8 PDZ interactions, so we sought to identify the mechanism supporting this transmission. We expressed GluA2_γ8 ΔNTD ΔPDZ on the AMPAR-null background, which failed to rescue AMPAR EPSCs (**Fig. 6c, 6g**), while NMDAR transmission was still intact (**Supplementary Figure 5a**).

As AMPARs at the CA1 synapse are mainly GluA1/2 heteromers^13^, we investigated whether these receptors were also able to anchor through their NTD alone. Co-expression of GluA1_γ8 and GluA2R_γ8 for rescue of AMPAR knockout gave non-rectifying responses on the cell surface (**Supplementary Figure 5c**) demonstrating heteromeric assembly, and synaptic transmission was successfully restored in these cells (**Figure 6d, 6h**). Removal of the γ8 PDZ-ligand from both subunits (GluA1_γ8 ΔPDZ & GluA2R_γ8 ΔPDZ) did not prevent surface trafficking, heteromeric assembly (**Supplementary Figure 5c**) or NMDAR mediated synaptic transmission (**Supplementary Figure 5b**). Critically however, while synaptic AMPAR EPSCs were reduced by this mutation, they were not abolished (**Fig. 6e, 6h**), suggesting that the GluA2 NTD can localise AMPARs at the synapse in the absence of other mechanisms. We tested this theory with two experiments.

First, we expressed γ8 PDZ-lacking heteromeric AMPARs with the GluA2 NTD swapped for that of GluA1 (denoted GluA2R^A1NTD^_γ8 ΔPDZ). Co-expression of GluA1_γ8 ΔPDZ and GluA2R^A1NTD^_γ8 ΔPDZ in AMPAR-null cells could not rescue synaptic AMPAR currents (**Fig. 6f, 6h**), while NMDAR currents remained unimpaired (**Supplementary Figure 5b**). Secondly, we transferred the GluA2 NTD onto GluA1_γ8 ΔPDZ, creating GluA1^A2NTD^_γ8 ΔPDZ (**Fig. 6i**). In contrast to GluA1_γ8 ΔPDZ (**Fig. 3b**), robust AMPAR transmission was recorded at GluA1^A2NTD^_γ8 ΔPDZ expressing synapses, demonstrating functional synaptic localisation (**Fig. 6i and Supplementary Figure 5d**). Together these data show that, while important for maintaining the full complement of synaptic AMPARs, γ8 PDZ interactions are not essential for CA1 synaptic transmission and work in concert with an anchoring mechanism mediated by the GluA2 NTD.

## Discussion

AMPAR trafficking defines the strength of excitatory synapses, and has therefore been investigated for decades^3,41^. Many molecular interactions have been reported, but there is incomplete synthesis of proposed models, which has muddied our understanding. Here, we demonstrate that while AMPAR CTD interactions play a modulatory role in shaping transmission, both intracellular interactions with the PSD through TARP γ8 and contacts mediated by the extracellular NTD control AMPAR transmission at CA1 synapses. These two interactions have unique roles: γ8 accumulates receptors at the PSD, while the NTD acts in a subunit-selective manner on a synaptic and subsynaptic level to allow functional transmission (**Fig. 7**).

**Fig. 7.**
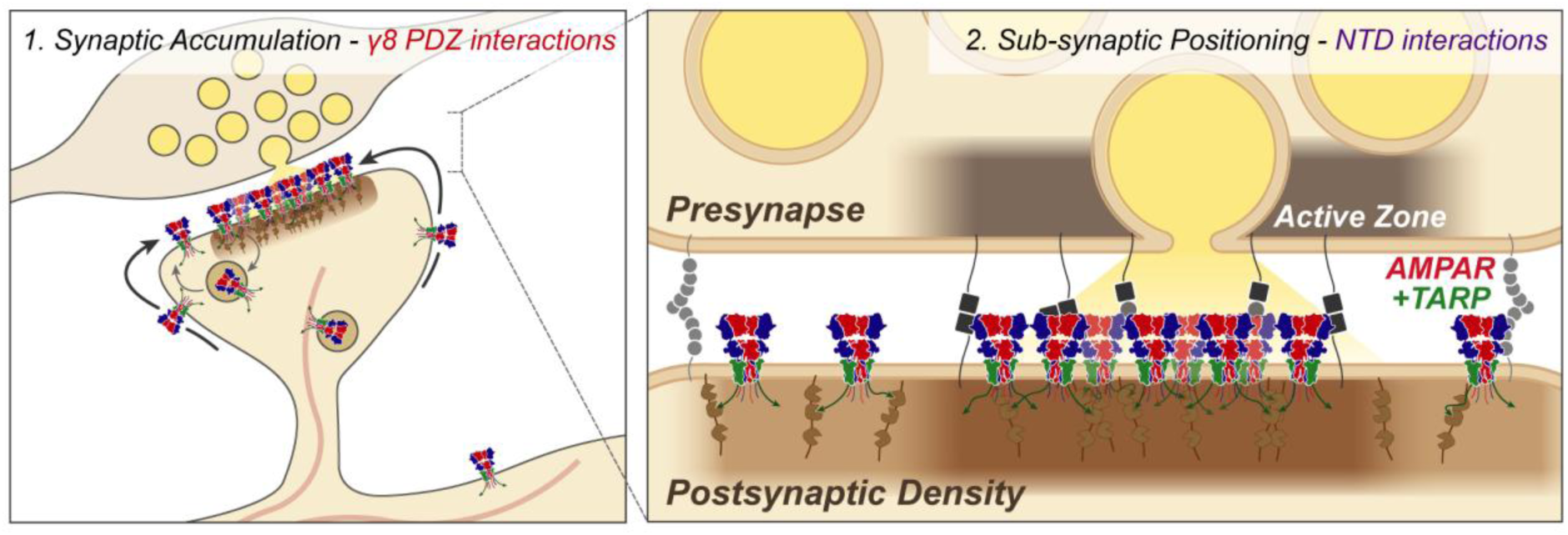
Proposed model outlining the differential role of TARP and NTD interactions on AMPAR CA1 synaptic anchoring. TARP PDZ and NTD interactions are critical for synaptic transmission. CTD interactions may influence receptor trafficking and recycling. TARP γ8 PDZ interactions drive accumulation of receptors at synaptic sites (left). NTD interactions may control subsynaptic receptor localization for contribution to functional transmission (right).

TARP PDZ interactions have become the principal model for AMPAR anchoring in recent years. First demonstrated at cerebellar granule cell synapses, where the PDZ-ligand of γ2 is essential for transmission^42^, this model was extended to hippocampal synapses^16,43,44^. However γ8, not γ2, is the predominant TARP in hippocampal neurons^14,15,45,46^. Mice with deletion of the γ8 PDZ-ligand show only a 30 % reduction in CA1 AMPAR transmission, while LTP remains intact^19^, questioning the requirement for PDZ interactions in hippocampal transmission. In contrast, Sheng et al.^36^ demonstrate complete dependence of GluA1_γ8 on PDZ_TARP_ interactions for both synaptic transmission and plasticity, concluding that all AMPAR transmission requires PDZ interactions with the PSD^36,47^. While we can directly replicate the PDZ-dependence of GluA1_γ8, this does not apply to all AMPARs. The GluA2 NTD acts as a minimal unit for synaptic anchoring even in the absence of γ8 PDZ ligands, and could support transmission in γ8 ΔPDZ mice^19^. While clearly an important mechanism in AMPAR synaptic localization, γ8 PDZ interactions are not the sole requirement for synaptic transmission. The role of other auxiliary subunits in synaptic anchoring, such as cornichons^14^, is excluded from our analysis, and remains to be determined.

The functional role of extracellular NTD interactions was reported only recently^27,28^. The dominant role of the GluA2 NTD described here, recapitulates the ‘synapto-sticky’ phenotype that we reported previously^27^, and will allow strict control over the contribution of GluA2-lacking, calcium permeable AMPARs^48,49^.

The mechanisms of NTD action are so far unclear. In contrast to γ8’s PDZ, NTD-removal does not prevent synaptic accumulation of receptors. The largely unaltered gross distribution of ΔNTD receptors contrasts starkly with their functional deficits, and suggests the domain has a subsynaptic influence on receptor organization at the synapse. Correspondingly, we observe changes in synaptic distribution of receptors on NTD removal, with ΔNTD receptors forming larger, less dense arrangements at the PSD. Within the PSD, AMPARs form nanoclusters^7,8^, and are positioned opposite vesicle release sites^10,50^. Surprisingly, nanocluster formation is not dependent on the NTD of either GluA1 or GluA2, suggesting that intracellular interactions with PSD-95 are sufficient for their formation^7^. PSD-95 forms nanoclusters that are trans-synaptically aligned with presynaptic release sites^7,51^.

It is intriguing that GluA1 ΔNTD receptors, which do not contribute to functional transmission, are still localized and clustered at synapses. NTD removal does alter the properties of these clusters suggesting a role for synaptic cleft interactors in AMPAR organization. There is emerging evidence that even modest changes in subsynaptic receptor distributions, such as clustering or trans-synaptic alignment, can strongly influence receptor activation^7,9^, or give rise to synaptic plasticity^52–54^. Therefore, the subtle rearrangements observed on NTD-deletion may cause substantial deficits in functional transmission. Given the non-uniform, protein-dense nature of the synaptic cleft^55,56^, a plethora of pre- or postsynaptic protein interaction partners would be able to tune AMPAR transmission through the NTD. Multiple reports demonstrate AMPAR-specific effects on deletion of trans-synaptic synaptic adhesion molecules such as Neurexins and LRRTMs^57,58^, and Pentraxin-dependent AMPAR clustering via the NTD has a clear role at other synapses in the brain ^59–61^.

Like others, we observe no essential role for the AMPAR CTD in constitutive receptor anchoring^25,26^, and no requirement for the GluA1 CTD PDZ ligand for synaptic transmission or potentiation^25,62,63^. Further investigation is required to assess their requirement for synaptic plasticity^26^. While not essential, we do observe some influence of CTDs, conferring dominant negative effects on NTD-deleted receptors. Specifically, the GluA2 CTD acts in basal transmission, and the GluA1 CTD in synaptic potentiation, echoing the subunit-specific AMPAR trafficking rules proposed to rely on CTD interactions^24^. While the CTD is not a critical player in AMPAR synaptic localization, it likely influences the trafficking and recycling of receptors^64^.

Changing the synaptic AMPAR content can control synaptic strength^65^, as demonstrated *in vivo* during behavioural tasks^66^. Our work proposes that γ8 PDZ and AMPAR NTD interactions are the major determinants of the postsynaptic receptor complement, with their interplay acting to trap and localize the required AMPAR content at each connection^5^. Plasticity-induced changes in the arrangement of the PSD are far better documented than those in the synaptic cleft^10,63,67^. Our data suggest that changes in the abundance or distribution of *extracellular* AMPAR binding slots may occur within the synaptic cleft, with the potential to serve as a mechanism for LTP. The identity and dynamics of these machineries, and the mechanisms of their pre-, post- or transsynaptic control is now of great importance to understanding information storage at synapses throughout the brain.

## Acknowledgements

The authors are very grateful to Andrew Penn for advice and discussions on surface receptor labelling in slice tissue, dissociated culture transfection, and for providing tdTomato and BirA^ER^ expression plasmids. This work would not have been possible without support from the Biological Services teams at both the Laboratory of Molecular Biology and Ares facilities. We are also very grateful to Nick Barry and Jerome Boulanger of the LMB Light Microscopy facility for support with confocal and STORM imaging and analysis, Junichi Takagi for providing scFv-Clasp expression constructs, Veronica Change for assistance with scFv-Clasp protein production, and Nejc Kejzar for assistance with cluster analysis. We would like to thank Teru Nakagawa and Ole Paulsen for critical reading of the manuscript and constructive feedback. This work was supported by grants from the Medical Research Council (MC_U105174197) and BBSRC (BB/N002113/1).

## Author Contributions

J.F.W.: performed and analysed electrophysiology and slice imaging experiments; A.P.: performed and analysed confocal and STORM imaging on cultured hippocampal neurons; H.H.: assisted with initial electrophysiology experiments; I.H.G. supervised the project and acquired all funding. All authors contributed to writing the manuscript.

## Declaration of Interests

The authors declare no competing interests.

## Materials and Methods

### Animals

All experimental procedures were performed in accordance with UK Home Office regulations and were licensed under the Animals (Scientific Procedures) Act of 1986 following local (AWERB) ethical approval. Tissue for slice or dissociated hippocampal cultures was harvested from an unascertained mixture of sexes at postnatal ages up to P8 either from C57BL/6JOla wild-type (RRID:MGI:3691859) or *Gria1-3*^*fl/fl*^ mice (reported in^27^), as indicated. To generate the *Gria1-3*^*fl/fl*^ line, mice with floxed loci at *Gria1, 2* and *3* genes [*Gria1*^*lox/lox*^ (RRID:IMSR_JAX:019012), *Gria2*^*lox/lox*^ (RRID:IMSR_EM:09212), *Gria3*^*lox/lox*^ (RRID:IMSR_EM:09215)] were interbred to produce mice homozygous for all floxed alleles (*Gria1*^*lox/lox*^; *Gria2*^*lox/lox*^; *Gria3*^*lox/lox*^, denoted *Gria1-3*^*fl/fl*^). All animals were housed with free access to food and water on a 12:12 hour light:dark cycle.

### Dissociated Hippocampal Culture

Hippocampal neurons from postnatal (P0-P1) C57BL/6JOla wild-type mice were prepared as described previously^68^. Briefly, mouse hippocampi were dissected in ice-cold HBSS (Ca^2+^ and Mg^2+^ free, Gibco, Cat.# 14175095) containing 0.11 mg ml^-1^ sodium pyruvate (Gibco, Cat.# 12539059), 0.1 % glucose and 10 mM HEPES (Gibco, Cat.# 15630056) and cells were dissociated with trypsin (0.25 % wt/vol, Gibco, Cat.# 15090-046). Neurons were plated onto poly-L-lysine coated glass coverslips (12 mm round coverslips, Corning, Cat.# 354085 for confocal imaging or 24 mm round coverslips #1.5, Glaswarenfabrik Karl Hecht GmbH & Co KG, Cat.# 1001/24_15 92100105080 for STORM imaging) in equilibrated culture medium, which contains 86.55 % Minimum Essential Medium (MEM) (Gibco, Cat.# 21090022), 10 % heat inactivated fetal bovine serum (Gibco Cat.# 11573397), 0.45 % glucose, 1 mM sodium pyruvate and 2 mM GlutaMax (Gibco, Cat.# 35050038). Cultures were kept at 37 °C and 5 % CO_2_ in equilibrated maintenance medium containing 96 % Neurobasal medium (Gibco, Cat.# 21103049), 2 % B-27 plus Supplement (Gibco, Cat.# A3582801) and 2 mM GlutaMax, until 21 days in vitro (DIV). Half of the medium was replaced every 4-7 days.

### Organotypic Slice Culture

Hippocampal organotypic slice cultures were prepared as previously described ^6969^. Hippocampi from P6-8 mice were isolated in ice-cold high-sucrose Gey’s balanced salt solution containing (in mM): 175 Sucrose, 50 NaCl, 2.5 KCl, 0.85 NaH_2_PO_4_, 0.66 KH_2_PO_4_, 2.7 NaHCO_3_, 0.28 MgSO_4_, 2 MgCl_2_, 0.5 CaCl_2_ and 25 glucose at pH 7.3. Hippocampi were cut into 300 μm thick slices using a McIlwain tissue chopper and cultured on Millicell cell culture inserts (Millipore Ltd) in equilibrated slice culture medium (37°C/5% CO2). Culture medium contained 78.5% MEM, 15% heat-inactivated horse serum, 2% B27 supplement, 2.5% 1 M HEPES, 1.5% 0.2 M GlutaMax supplement, 0.5% 0.05 M ascorbic acid, with additional 1 mM CaCl_2_ and 1 mM MgSO_4_ (all from Thermo Fisher Scientific; Waltham, MA). Medium was refreshed every 3–4 days. Cultures were transfected at 5-10 DIV by single-cell electroporation (SCE) and recordings were performed 4-5 days after transfection.

### DNA Constructs

All constructs were created by *in vivo* assembly cloning unless otherwise stated ^7070^ using the *Rattus norvegicus* coding sequence. GluA1 (flip isoform; Uniprot P19490) or GluA2 (<u>Q</u>/R edited where specified, R/G edited, flip isoform; Uniprot P19491) variants used for electrophysiology and slice tissue imaging were expressed from the pRK5 vector. ΔNTD constructs were as reported previously^27^. The NTD coding sequence (GluA1: Ala1 - Ala373, GluA2: Val1 - Thr377) was replaced by a c-myc epitope sequence, leaving the entire NTD-LBD linker sequence intact. GluA2 NSF mutation was achieved by Asn830Ala and Pro831Ala mutations, PDZ by addition of a C-terminal Tyr residue (Tyr863) and CTD-null by all aforementioned mutations plus Ser842Ala. For GluA1/2 intracellular domain swap, GluA2 Thr553 to Thr568 (loop 1) and Ala820 to Iso862 (CTD) were replaced by GluA1 sequence from Ser549 to Ser564 (loop 1) and Ser816 to Leu889 (CTD). GluA1 PDZ mutation was achieved by Thr887Ala mutation, and was combined with Ser831Ala and Ser845Ala mutations to generate the CTD-null construct. N-terminal GFP tagging was achieved by insertion of the EGFP coding sequence between GluA1 Phe3 and Pro4 residues, flanked by Ala-Arg and Ala-Ser linker residues respectively (for replication of ^24^).

TARP tandem constructs were produced by splicing of the TARP γ8 coding sequence (Uniprot Q8VHW2) in frame with AMPAR subunits, separated by a Gly-Ser-Gly-Ser-Gly linker sequence (see ^13^). PDZ-deletion was achieved by removal of the γ8 terminal Tyr-Tyr-Pro-Val sequence (Tyr418 - Val421). GluA1/2 NTD swaps were performed as previously reported^27^, and involved exchanging both NTD and NTD-LBD linker sequence between subunits (corresponding to, GluA1: Ala1 - Thr390, GluA2: Val1 - Thr394). pN1-EGFP (Clontech, 6085-1) was used for cell visualisation.

For surface biotinylation, GluA1 was tagged at the N-terminus with the acceptor peptide sequence prior to Ala1 of GluA1 coding sequence (Insertion of Ala-*Gly-Leu-Asn-Asp-Iso-Phe-Glu-Ala-Gln-Lys-Iso-Glu-Trp-His-Glu*-Gly sequence, AP-tag in italics). GluA1 ΔNTD was tagged by replacing the c-myc epitope with the AP-tag sequence. The Lys residue in this sequence is modified by biotin conjugation (see ^71^). ER-localised biotin ligase (BirA), and pC1-tdTomato were gifts from Andrew Penn, and were used to create the pC1-tdTomato-IRES-BirA^ER^ construct.

For confocal and 3D STORM imaging of cultured hippocampal neurons, N-terminally HA-tagged AMPAR constructs were expressed from the doxycycline inducible pcDNA4/TO vector (Invitrogen Cat. No: V102020, including pcDNA™6/TR for expression of the tetracycline repressor (TR) protein). The HA-epitope was inserted after Ala1 for GluA1 full-length (flip isoform) and after Val1 for GluA2 full-length (<u>**Q**</u>/R unedited, R/<u>**G**</u> edited, flip isoform) (Insertion for GluA1: Ala-*Tyr-Pro-Tyr-Asp-Val-Pro-Asp-Tyr-Ala*, Insertion for GluA2: Val-*Tyr-Pro-Tyr-Asp-Val-Pro-Asp-Ala*, HA-tag in italics). HA-tagged GluA1 and GluA2 ΔNTD constructs are equivalent to those previously reported, with the c-myc epitope replaces by the HA-tag sequence. pN1-EGFP (Clontech, 6085-1) was used for cell visualisation.

Constructs for 12CA5 scFv-Clasp anti-HA (CA5-VH(X3)-Mst1_pET11c (#5019), heavy chain, and CA5-VL-Mst1(Y3)_pET11c (#5020), light chain) were kindly provided by Junichi Takagi^37^. The constructs were subcloned using restriction digestion into the pHL-sec vector containing a C-terminal 6xHis-tag (Addgene # 99845). Restriction sites for subcloning were introduced by PCR at the 5’ and 3’ ends of scFv-Clasp heavy and light chains (light chain: forward primer 5’-TAGTAGTAACCGGTCATATGGAAGTGA-AGTTGGTTGAATCTG-3’, inserting AgeI site, reverse primer 5’-TAGTAGTAGGTAC-CCTTAGCCTCTATGGCATCCAGG-3’, adding KpnI site; heavy chain: forward primer 5’-TAGTAGTAACCGGTATGGACATCGAATTGACTCAGTCTCC-3’, inserting AgeI site, reverse primer 5’-TAGTAGTATGTACATTACTTAGCCTCTATGGCATCCA-GG-3’, adding Bsp1407I site and removing stop codon). The light chain inset was cloned into the pHL-sec vector using AgeI and KpnI restriction sites, keeping a stop codon before the His-tag. The heavy chain sequence was introduced using AgeI and Bsp1407I (inset)/Acc65I (vector) sites, in frame with the His-tag.

### Transfection of neuronal cultures

Cultured hippocampal neurons were transfected at DIV 6 with plasmids expressing doxycycline-inducible, HA-tagged GluA1 or GluA2 receptors. AMPAR-expressing constructs were co-transfected with pcDNA6/TR and EGFP-pN1 at a 1:1.57:0.29 ratio using Ca_2_PO_4_ precipitation. 2 µg (12 mm coverslips) or 4 µg (24 mm coverslips) of DNA were added to 250 mM CaCl_2_ in nuclease free water, mixed at a 1:1 ratio with 2x HEPES buffered saline solution solution (50 mM HEPES, 280 mM NaCl, 1.5 mM Na_2_HPO_4_ in nuclease free water, pH 7.05) to a final volume of 100 (12 mm coverslips) or 200 µl (24 mm coverslips) and incubated for 20 minutes at room temperature. Coverslips containing neurons were moved to fresh plates with conditioned maintenance medium supplemented with 1 mM Kynurenic acid (Sigma, Cat.# K3375, 0.5 ml/12-well for 12 mm and 1 ml/6-well for 24 mm coverslips). 100-200 µl transfection mix were added to each 12- or 6-well, respectively, and incubated for 1.5-2 h at 37 °C and 5 % CO_2_. Subsequently, coverslips were washed in maintenance medium conditioned at 37 °C and 10 % CO_2_ (1 ml/12-well and 2 ml/6-well) for 20 min at 37 °C and 5 % CO_2_ to dissolve precipitates and eventually moved back to their original culture wells. Exogenous AMPAR expression was induced at DIV 18 using 7.5 µg/ml doxycycline (Merck, Cat.# D3447) and neurons were harvested at DIV 21 for imaging.

### Neonatal Viral Injection

*In vivo* viral injection was performed according to ^72^. P1 neonatal mice were anaesthetised using isoflurane, bilaterally injected using a pulled glass needle in the hippocampal area with 0.5 μL AAV-hSyn-Cre-EGFP at 3 × 10^12^ GC ml^-1^ (Addgene #105540, virus produced by U-Penn Vector Core), and returned to the parent cage until P7, when tissue was harvested for organotypic culture.

### Single-cell electroporation (SCE)

CA1 cells of hippocampal organotypic slices were transfected using an adapted version of the single-cell electroporation method described in ^73^. DNA plasmids were diluted to 33 ng/μL with potassium-based intracellular solution and the mixture was back-filled into borosilicate microelectrode pipettes. Slices were submerged in HEPES-based artificial cerebrospinal fluid (aCSF) containing (in mM): 140 NaCl, 3.5 KCl, 1 MgCl_2_, 2.5 CaCl_2_, 10 HEPES, 10 Glucose, 1 sodium pyruvate, 2 NaHCO_3_, at pH 7.3. Plasmids were introduced into individual cells by the application of a short burst of current pulses (60 pulses at 200 Hz) while in cell-attached mode. To visualize transfected cells, pN1-EGFP or pC1-tdTomato (Clontech; Mountain View, CA) were mixed with AMPAR-expressing plasmids at a base pair ratio of 1:7. For heteromeric receptor transfection, GluA1 and GluA2 plasmids were transfected at a 1:2 ratio. In tCaMKII experiments, the ratio between tCaMKII-EGFP and AMPAR-expressing plasmids was 1:1. For rescue of AMPAR-null cells, SCE was used to target neurons strongly labelled with nuclear EGFP fluorescence from *in vivo* viral transduction. For slice imaging experiments, AP-AMPAR and tdTomato-IRES-BirA^ER^ plasmids were transfected at a ratio of 3:1.

### Electrophysiology

Transfected hippocampal slice cultures were submerged in aCSF containing (in mM): 125 NaCl, 2.5 KCl, 1.25 NaH_2_PO_4_, 25 NaHCO_3_, 10 glucose, 1 sodium pyruvate, 4 CaCl_2_, 4 MgCl_2_ and 0.001 SR-95531 (Tocris Cat# 1262) at pH 7.3 and saturated with 95% O_2_/5% CO_2_. 100 μM D-APV (Tocris Cat# 0106, HelloBio Cat# HB0225) was used to isolate AMPAR currents for rectification index recordings. 2 μM 2-chloroadenosine (Tocris Cat#3136) was added to aCSF to dampen epileptiform activity. 3–6 MΩ borosilicate pipettes were filled with intracellular solution containing (in mM): 135 CH_3_SO_3_H, 135 CsOH, 4 NaCl, 2 MgCl_2_, 10 HEPES, 4 Na_2_-ATP, 0.4 Na-GTP, 0.15 spermine, 0.6 EGTA, 0.1 CaCl_2_, at pH 7.25. All synaptic recordings were performed using dual whole-cell patch clamp involving simultaneous recording from a neighboring pair of GFP positive and negative cells for exogenous expression experiments. For AMPAR rescue experiments, dual recordings were performed from one EGFP (Cre) and tdTomato (SCE rescue) positive cell, and a neighbouring non-fluorescent cell. EPSCs were evoked by single simulations of Schaffer collaterals in the stratum radiatum of CA1 using a monopolar glass electrode, filled with aCSF at 0.2 Hz. Recordings were collected using a Multiclamp 700B amplifier (Axon Instruments), digitised using a Digidata 1440A interface (Axon Instruments) and recorded using Clampex (pClamp10).

Somatic AMPAR responses were recorded by pulling outside-out patches from fluorescence positive or negative CA1 cell bodies and patches were subjected to fast-exchange perfusion using a two-barrel theta glass tube controlled by a piezoelectric translator (Physik Instrumente). Theta barrels solutions were HEPES-based aCSF (see SCE) containing 100 μM cyclothiazide (Tocris Cat# 0713) and 100 μM D-APV, either with or without 1 mM L-glutamate. In voltage-clamp mode, a 500 ms holding potential ramp from −100 mV to +100 mV was applied to patches (see Figure S1).

### Electrophysiology data analysis

Data were analysed using Clampfit (pClamp 10 - Molecular Devices). Synaptic stimulations across at least 20 sweeps were averaged, and peak amplitudes were measured relative to baseline preceding the simulation artifact. For rectification index calculation, only recordings with temporal alignment of peak amplitudes between negative (−60 mV) and positive (+40 mV) holding potentials were included in analysis. Rectification index was calculated from peak current amplitudes at -60, 0 and +40 mV holding potentials as follows:

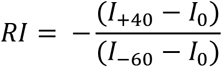

AMPAR peak amplitudes were measured from -60 mV holding potential recordings. NMDAR synaptic currents were measured from +40 mV holding potential recordings, and represent the amplitude 100 ms after the stimulus artifact, to prevent AMPAR current contamination. Normalisation of dual recording peak amplitudes was performed by division of all values across the cell pair by the mean amplitude of all control cell responses. Where normalized summary graphs are depicted without individual points, all amplitude values are presented as scatter graphs in either the Main Text Figures or Supplementary Information. Somatic response rectification indices are calculated from current amplitudes from membrane potential ramps (−100 to +100 mV). Recordings in the absence of glutamate were subtracted from those in the presence of glutamate for each patch. All recordings during which the series resistance varied by more than 20% or exceeded 20 MΩ were discarded. Membrane potentials are given without correction for liquid junction potentials. All data are pooled recordings from at least 2 individual preparations, except surface rectification recordings which are form 1 preparation.

### Generation of scFv-Clasp against HA

HEK293S GnTI– cells (ATCC, Cat.# CRL-3022, RRID:CVCL_A785) were transiently transfected with heavy (CA5-VH(X3)-His_pHL_sec) and light chains (CA5-VL(X3)_pHL_sec) at a 1:1 ratio using polyethyleneimine (PEI, 1 mg/ml, MW 40,000, Polysciences Cat.# 24765) transfection reagent. A mixture of DNA (0.5 mg) and PEI (1 mg) diluted in 20 ml serum free culture medium (Dulbecco’s Modified Eagle Medium (DMEM), High glucose, GlutaMax supplement, pyruvate, Gibco, Cat.# 31966-021) was added per 250 ml of culture volume. Cells were maintained in culture medium (DMEM, as described) supplemented with 2 % FBS, 2 mM Glutamax, and 1 % MEM Non-Essential Amino Acids Solution, Gibco, Cat.# 11140050) at 30 °C for 7 days after transfection (see ^74^). Cleared cell supernatant, containing secreted scFv-Clasp, was concentrated using an Akta Flux and stored over night at 4 °C in 20 mM Tris HCl pH 8, before addition of 4 mM imidazole and incubation on a rotor (111 RPM) with TALON metal affinity resin (Takara, Cat.# 635503, 4 ml slurry for ∼300 ml concentrated supernatant) at 16 °C, for 4 h the following day. After collection, beads were washed with 20 mM Tris/PBS pH 8, non-specific proteins were eluted with 10 mM imidazole in 20 mM Tris-HCl and 500 mM NaCl, pH 8, followed by protein elution with 250 mM imidazole in 20 mM Tris-HCl, 500 mM NaCl, pH 8. Fractions containing scFv-clasp were pooled, concentrated and further purified by size-exclusion chromatography on a Superdex 200 26/60 column L (GE Healthcare, Cat.# 28-9893-36). Purified scFv-Clasp were flash frozen in aliquots (in 10 mM HEPES, 150 mM NaCl, pH 7.4) and stored at -80 °C until use.

### scFv-Clasp fluorescent conjugation

scFv-Clasp (∼1 mg/ml) was adjusted to pH 8.3 with 1M NaHCO_3_, incubated either with Alexa Fluor (AF) 647 NHS ester (Invitrogen, Cat.# A20106) or CF568 NHS ester (Sigma-Aldrich, Cat.# SCJ4600027), according to manufacturer’s instructions, for 1.5-2 h while shaking at room temperature in the dark. Subsequently, the solution was centrifuged for 10 min at 10,000 g at 4 °C. If necessary, the volume was adjusted to 500 µl with PBS, and loaded on a equilibrated G-25 column (Column PD Minitrap G-25, GE Healthcare, Cat.# 28-9180-07). Once the solution fully entered the column, conjugated scFv-Clasp was eluted with 1 ml PBS. To remove any residual unbound dye, the eluate was washed 3 times with PBS in an Amicon Ultra-4 Centrifugal Filter Unit (NMWL 3 kDa, Millipore, UFC800324) and finally concentrated to < 250 µl. The degree of labelling, assessed photometrically, was 5.5 for AF647-scFv-Clasp and 2 for CF568-scFv-Clasp.

### Surface labelling of biotinylated receptors in slice tissue

Surface labelling of biotinylated AMPARs in slice tissue was performed as described in ^5^. 20 μM biotin (Sigma-Aldrich Cat# B4501) was added to the culture medium after SCE transfection. 4 days later, slices were washed in HEPES-based aCSF (see SCE) and excess biotin was removed by dialysis in HEPES-based aCSF against 0.25 ml 2 mg/ml avidin Texas red (Thermo Fisher Scientific Cat# A820) through a Slide-A-Lyzer MINI (3500 molecular weight cut off, Thermo Fisher) for 45 mins. Slices were then incubated for 45 min at room temperature in HEPES-based aCSF containing 120 nM Streptavidin-AF647 (Invitrogen). Slices were washed 3 times over 45 mins before fixation in phosphate-buffered saline (PBS) containing 4 % paraformaldehyde and 4 % sucrose (PFA, Sigma-Aldrich, Cat.# P6148) at 4 °C overnight. Slices were washed and stored in PBS until imaging. For imaging, slices were inverted on a glass-coverslip and imaged in PBS using a 63x/1.4NA oil-immersion objective on a commercial Leica TCS SP8 confocal microscope using LAS X software. Z-stacks of whole cells or whole stratum radiatum dendrites were taken, with EGFP, tdTomato and AF647 excited with 478 nm, 554 nm and 653 nm laser lines, with equivalent power settings across experimental conditions.

### Immunostaining

Cultured hippocampal neurons were live-labelled at DIV 21 with 50 nM scFv-Clasp either conjugated to Alexa647 (confocal imaging) or CF568 (STORM imaging) at room temperature for 20 min in aCSF (in mM: 120 NaCl, 5 KCl, 1.2 MgCl_2_*6H_2_O, 2 CaCl_2_, 25 HEPES, 30 glucose, pH set to 7.3-7.4 with 2M NaOH). Neurons were briefly washed 5 times over 10 min in aCSF and PBS and fixed in 4 % PFA, 4 % sucrose in PBS for 10 min at room temperature. Subsequently cells were washed again, permeabilised with 0.1 % Triton X-100 (Fisher Bioreagents, Cat.# BP151-500) and treated with blocking solution containing 1 % bovine serum albumin (Fisher Bioreagents, Cat.# BP1605-100) and 10 % normal goat serum (Sigma-Aldrich, Cat.# G9023) in PBS. Neurons were incubated sequentially with primary and secondary antibody solutions prepared in PBS supplemented with 1 % BSA and 10 % normal goat serum for 2 h at room temperature, and washed in PBS after each incubation. For 3D STORM imaging a second fixation was performed after incubation with the secondary antibody and coverslips were stored in PBS at 4 °C until imaging. For confocal imaging, coverslips were mounted in ProLong Diamond antifade Mountant (Invitrogen, Cat.# P36961) and left to cure for 48 h at room temperature before imaging.

### Antibodies

The following antibodies were used in this study: rabbit anti-HA (Sigma-Aldrich, Cat.# H6908, RRID:AB_260070, 1:200, confocal imaging), rabbit anti-bassoon (Synaptic Systems Cat# 141 003, RRID:AB_887697, 1:200, STORM imaging) goat anti-rabbit IgG AF568 (Thermo Fisher Scientific Cat# A-11036, RRID:AB_10563566, 1:200, confocal imaging), goat anti-rabbit IgG AF647 (Thermo Fisher Scientific Cat# A-21245, RRID:AB_2535813, 1:200, STORM imaging).

### Confocal Imaging

Images were acquired with a Leica TCS SP8 confocal inverted microscope controlled by Leica Application Suite X (LAS X) software and equipped with a tuneable pulsed white light laser and hybrid detectors. For quantification of AMPAR expression levels and surface distribution in cultured hippocampal neurons, z-stacks of whole dendrite segments were acquired using a 63x/1.4NA oil-immersion objective. EGFP, Alexa Fluor 568 and Alexa 647 were excited at 488 nm, 561 nm and 633 nm, respectively. All conditions were acquired with the same settings across different preparations.

### Confocal Imaging Analysis

For AMPAR surface distribution in tissue, images were analysed in FIJI. Dendritic rejoins were z-projected (sum) before median filtering using a 1 pixel radius filter. Line profiles were performed on all spines that were suitably perpendicular to the dendritic shaft within each field of view, from tdTomato fluorescence. Grey values for surface receptor fluorescence were measured along a line of 300 μm thickness from raw images. Line profiles were normalised to the maximal dendritic fluorescence, and spine enrichment is quantified as the peak spine fluorescence after normalisation. Quantifications in figures 3e and 3f are from 2 and 1 preparations respectively.

Confocal images of cultured hippocampal neurons were z-projected (maximal intensity) and a median filter (radius = 1) was applied. For quantification of expression levels, masks of regions of interests, generated from thresholded composite images of permeabilized and surface HA stains were applied to each channel and integrated densities were determined.

Thresholds and filters were applied to the entire image and same settings were used across all conditions. For surface distribution, line scans across spines perpendicular to the dendritic shaft were performed based on EGFP fluorescence. Spine enrichment was calculated as peak spine fluorescence over peak dendrite fluorescence. Fluorescence intensity was normalized to maximal dendritic intensity for representative line scans in Supplementary Figure 4c-d. Same datasets were used for quantification of expression levels and surface distribution, two images with the lowest surface expression were excluded for analyzing GluA2 ΔNTD spine enrichment. Images shown in Figure 4b-c were processed as described above. Where needed, contrast adjustments were performed uniformly across the whole image, with same settings applied to full-length and respective ΔNTD AMPAR example images. Images were analysed and pseudo-coloured in Fiji (ImageJ).

### STORM Imaging

Samples were incubated with Tetraspeck Microspheres (100 nm, Invitrogen, Cat.# T7279) for image registration and mounted in imaging buffer containing 5 % Glucose (w/v), 100 mM Mercaptoethylamine (Sigma-Aldrich, Cat.# 30070), 0.8 mg/ml glucose oxidase (Sigma-Aldrich, Cat.# G2133), 50 µg/ml catalase (Sigma-Aldrich, Cat.# C30) in PBS in an Attofluor Cell Chamber (Invitrogen, Cat. # A7816). The imaging buffer was freshly prepared for each coverslip. The imaging chamber was filled with buffer to full capacity and sealed by an additional coverslip on top to minimize air-induced oxidation of the sample. 3D STORM images were acquired using a Nikon Eclipse TI-E N-STORM inverted microscope equipped with an Apochromat TIRF 100x/1.49 NA oil immersion objective, 405nm, 488nm, 561nm and 647nm lasers and an EMCCD camera (iXon Ultra DU897, Andor). The illumination intensity at the sample was ∼ 2KW/cm^2^. 3D localisation was achieved using the astigmatism method ^75^. Conventional fluorescence images for different channels were acquired first. Prior to STORM image collection, the field of view was illuminated with the activating laser at a low incident angle to deactivate out-of-focus fluorophores. For STORM data acquisition, samples were illuminated in pseudo-TIRF mode with the incident angle close to the critical angle with either the 647 nm (AF647) or the 568 nm (CF568) lasers using 100 % intensity. The 405 nm laser was used at 2 % to reactivate AF647 fluorophores to maintain adequate localisation density. 50,000 frames at 50 Hz, were collected for each channel sequentially, the axial focal plane was maintained by using the Perfect Focus System during image acquisition.

### STORM imaging analysis

Single molecule identification was performed with NIS Elements Advanced Research software (Nikon) using default settings for peak identification with minimum intensity (height) set to 1000. Identical settings were used for every image. Lateral drift correction was performed using in-built cross-correlation between subsets for the frame sequence. Molecules outside the Z calibration range, appearing in more than 5 consecutive frames, with fewer than 500 photons and a localization precision greater than 20 nm, were discarded. Tetraspeck Microspheres were used for image registration. Individual localizations in the reconstructed images (Figure 4e-h) are shown as a normalized Gaussian with the width corresponding to the localization uncertainty. Images were thresholded uniformly, with the same settings applied to full-length versus respective ΔNTD AMPAR images. For visualization of dendrite outlines, conventional fluorescence images of co-transfected EGFP, were smoothened, rendered semi-transparent and overlaid onto 3D STORM images using Adobe Photoshop. For density plots (Figure 5a), synaptic AMPAR clusters were reconstructed as scatter plots in MATLAB (Mathworks) and colour coded based on their local density (number of points (60) within search radius defined as 99^th^ percentile of shortest pair-wise distances). As control, observed localisations were randomly re-distributed within the original cluster boundaries.

Quantitative analysis on imported molecule lists was performed in MATLAB using custom scripts. STORM images were reconstructed as scatter plots and isolated synapses, defined as opposing post-synaptic AMPAR and pre-synaptic Bassoon clusters were selected for analysis. Only synaptic clusters with an unambiguous orientation were selected and rotated with the synaptic cleft perpendicular to the z-axis. Noise points (i.e. localisations on the outer edges of synaptic clusters) and nanoclusters were identified using the OPTICS (Ordering points to identify the clustering structure) algorithm which offers superior robustness when working with datasets of different densities compared to other methods such as DBSCAN^38–40^. OPTICS scripts for calculating the reachability distances and subsequent cluster extraction were adapted from Michal Daszykowski (http://chemometria.us.edu.pl/index.php?goto=downloads) and Alex Kendall (https://github.com/alexgkendall/OPTICS_Clustering), respectively, with minor modifications for increased robustness. To generate ‘*reachability plots’* representing the hierarchical structure of clusters, points were linearly ordered based on their spatial relationship with minimum number of points (MinPts) set to 30 (noise identification) or 25 (nanocluster detection). Each channel was analysed separately. To reduce bias, results from 100 repeated runs, each starting from a different localisation, were averaged for individual clusters. Averaged reachability distances greater than the 90^th^ percentile were used as the cut-off to define noise points. Subsequently, synaptic areas were analysed using the in-built MATLAB function *‘boundary’* and density was calculated as number of localisations per area. Clusters with a synaptic area > 0.6 µm^2^ were excluded from analysis. For nanocluster quantification, the smallest detected clusters, if present, containing a minimum number of 25 points were extracted from individual reachability plots as described in (^38^, ξ set to 0.02) using an adaptation of the aforementioned MATLAB script by Alex Kendall. Due to variability in cluster edge detection, only the localisations that were detected as part of a putative cluster in over 70 % of the iterations were considered to constitute to a nanocluster. Furthermore, only nanoclusters with a ratio of averaged reachability distance to overall reachability distance greater than 0.8, more than 25 localisations and a density (localisations/area) greater than 3×10^3^ were considered for analysis. Nanocluster detection criteria were determined based on randomised datasets, which were generated by subjecting the observed localisations in each AMPAR synaptic cluster to random re-distribution within the measured cluster boundaries. 10 different randomisations were performed for each dataset, analysed as described above, giving an estimated nanocluster detection rate below 0.5 % within the randomly re-distributed localisations dataset. Nanocluster volumes were determined using MATLAB inbuilt functions ‘*alphaShape*’ and density was calculated as number of localisations per volume. Diameters were defined as the longest axis across the nanocluster area, determined by the function ‘*boundary*’.

### Statistical analysis

Statistical testing of rectification index dual recordings was performed using a two-tailed paired t-test, as data is normally distributed paired data. Testing of amplitude comparisons from dual recordings was performed using a two-sided Wilcoxon matched-pairs signed rank test on data prior to normalization, as peak amplitudes are non-parametric. All bars represent mean value, ± standard error of the mean (SEM), unless stated otherwise. For surface receptor imaging in tissue, box and whiskers represent mean, upper and lower quartiles (box), and min to max values (whiskers). Samples were statistically tested using a Mann-Whitney test as data is non-parametric. The expression levels (Supplementary Figure 4a-b) and synaptic properties such as areas, densities and localisations (Figures 4i-j, Supplementary Figure 4e) of ΔNTD AMPARs and corresponding full-length receptors in cultured hippocampal neurons were compared using unpaired t-test as data was normally distributed; corresponding data is shown as mean ± S.E.M. For nonparametric data such as surface distribution (Figure 4d), and nanocluster properties as number, diameter, localization and densities (Figure 5b-e) statistical significance was assessed using Mann-Whitney test and data is shown as median ± 95 % confidence interval (CI). Relative frequency distributions (Supplementary Figure 4f-g) were analysed using Kolmogorov-Smirnov test. Slopes of linear regressions (Figure 4k-l) were compared with F-test. Sample sizes and biological replicates are given in the figure legends. Statistical analysis was performed in GraphPad Prism 8. Throughout the manuscript, * indicates p < 0.05, ** p < 0.01, *** p < 0.001 and ^ns^ specifies no significance.

## Data Availability

Data supporting the findings of these study are available from the corresponding author upon reasonable request.

## Code Availability

The image analysis code supporting the current study is available from the corresponding author on request. Original OPTICS MATLAB scripts by Michal Daszykowski and Alex Kendall can be found at http://chemometria.us.edu.pl/index.php?goto=downloads, and https://github.com/alexgkendall/OPTICS_Clustering, respectively.

## Notes

### Competing Interest Statement

The authors have declared no competing interest.

