## Supplementary Information for "AMPA receptor anchoring at CA1 synapses is determined by an interplay of N-terminal domain and TARP γ8 interactions"

**Containing:**

Supplementary Tables 1-2

Supplementary Figures 1-5

**Supplementary Table 1: Dual Cell Synaptic Current Properties - GluA2Q data**

| | Rectification Index | | | |
|---|---|---|---|---|
| **Construct** | **Ctrl** | **Transf.** | **n (pairs)** | **p*** |
| GluA2Q | 0.61 ± 0.05 | 0.22 ± 0.03 | 6 | 0.0009 |
| GluA2Q ΔNSF | 0.62 ± 0.04 | 0.19 ± 0.028 | 11 | <0.0001 |
| GluA2Q ΔPDZ | 0.60 ± 0.08 | 0.24 ± 0.04 | 8 | 0.0008 |
| GluA2Q A1ICD | 0.49 ± 0.05 | 0.17 ± 0.03 | 12 | <0.0001 |
| GluA2Q CTD-null | 0.62 ± 0.04 | 0.19 ± 0.03 | 15 | <0.0001 |
| GluA2Q ΔNTD | 0.60 ± 0.04 | 0.21 ± 0.03 | 9 | <0.0001 |
| GluA2Q ΔNTD ΔNSF | 0.56 ± 0.05 | 0.38 ± 0.03 | 10 | 0.013 |
| GluA2Q ΔNTD ΔPDZ | 0.57 ± 0.04 | 0.27 ± 0.03 | 12 | <0.0001 |
| GluA2Q ΔNTD A1ICD | 0.56 ± 0.05 | 0.41 ± 0.03 | 9 | 0.018 |
| GluA2Q ΔNTD CTD-null | 0.61 ± 0.02 | 0.15 ± 0.02 | 15 | <0.0001 |

| | AMPAR EPSC Amplitudes (pA) | | | |
|---|---|---|---|---|
| **Construct** | **Ctrl** | **Transf.** | **n (pairs)** | **p*** |
| GluA2Q | 37.4 ± 10.2 | 54.8 ± 10.2 | 8 | 0.0078 |
| GluA2Q ΔNSF | 34.8 ± 3.9 | 48.2 ± 7.5 | 11 | 0.024 |
| GluA2Q ΔPDZ | 23.1 ± 2.9 | 34.7 ± 5.3 | 9 | 0.0039 |
| GluA2Q A1ICD | 39.1 ± 8.0 | 54.6 ± 9.1 | 12 | 0.043 |
| GluA2Q CTD-null | 53.1 ± 6.3 | 65.7 ± 8.5 | 20 | 0.030 |
| GluA2Q ΔNTD | 40.6 ± 4.1 | 21.8 ± 2.2 | 14 | 0.0001 |
| GluA2Q ΔNTD ΔNSF | 37.6 ± 3.2 | 29.4 ± 4.5 | 11 | 0.024 |
| GluA2Q ΔNTD ΔPDZ | 91.4 ± 15.5 | 65.8 ± 11.0 | 12 | 0.016 |
| GluA2Q ΔNTD A1ICD | 38.6 ± 5.0 | 31.7 ± 6.1 | 9 | 0.049 |
| GluA2Q ΔNTD CTD-null | 50.0 ± 4.5 | 30.2 ± 3.6 | 15 | 0.0002 |

** Statistical analysis - Rectification Index: two-tailed paired t-test; EPSC ampltiudes: Wilcoxon matched-pairs signed rank test.*

**Supplementary Table 2: Dual Cell Synaptic Current Properties - GluA1 data**

| | Rectification Index | | | |
|---|---|---|---|---|
| **Construct** | **Ctrl** | **Transf.** | **n (pairs)** | **p*** |
| GluA1 | 0.53 ± 0.03 | 0.37 ± 0.03 | 7 | 0.0058 |
| GluA1 ΔPDZ | 0.60 ± 0.04 | 0.38 ± 0.04 | 13 | <0.0001 |
| GluA1 ΔPDZ +tCKII | 0.58 ± 0.06 | 0.41 ± 0.05 | 12 | 0.012 |
| GluA1 CTD-null | 0.58 ± 0.04 | 0.45 ± 0.04 | 17 | 0.0016 |
| GluA1 + TTX incubation | 0.64 ± 0.07 | 0.31 ± 0.03 | 7 | 0.0082 |
| GFP-GluA1 | 0.60 ± 0.04 | 0.54 ± 0.04 | 16 | 0.086 |
| GluA1 ΔNTD | 0.57 ± 0.07 | 0.53 ± 0.05 | 9 | 0.54 |
| GluA1 ΔNTD ΔPDZ | 0.52 ± 0.05 | 0.49 ± 0.04 | 9 | 0.41 |
| GluA1 ΔNTD +tCKII | 0.56 ± 0.05 | 0.45 ± 0.05 | 10 | 0.0083 |
| GluA1 ΔNTD ΔPDZ + tCKII | 0.63 ± 0.03 | 0.60 ± 0.05 | 8 | 0.70 |

| | AMPAR EPSC Amplitudes (pA) | | | |
|---|---|---|---|---|
| **Construct** | **Ctrl** | **Transf.** | **n (pairs)** | **p*** |
| GluA1 | 24.5 ± 1.4 | 19.6 ± 1.8 | 7 | 0.078 |
| GluA1 ΔPDZ | 60.8 ± 7.2 | 56.4 ± 8.2 pA | 14 | 0.30 |
| GluA1 ΔPDZ +tCKII | 50.8 ± 9.0 | 106.2 ± 16.0 | 8 | 0.0078 |
| GluA1 CTD-null | 46.8 ± 5.8 | 37.9 ± 3.8 | 17 | 0.38 |
| GluA1 + TTX incubation | 72.3 ± 12.7 | 52.2 ± 5.6 | 8 | 0.15 |
| GFP-GluA1 | 65.3 ± 5.7 | 49.8 ± 5.4 | 13 | 0.13 |
| GluA1 ΔNTD | 29.5 ± 4.9 | 19.1 ± 2.3 | 9 | 0.055 |
| GluA1 ΔNTD ΔPDZ | 59.2 ± 6.1 | 47.5 ± 4.4 | 9 | 0.30 |
| GluA1 ΔNTD +tCKII | 49.9 ± 6.8 | 43.1 ± 12.0 | 9 | 0.13 |
| GluA1 ΔNTD ΔPDZ + tCKII | 21.7 ± 2.7 | 38.8 ± 4.7 | 16 | 0.0016 |

** Statistical analysis - Rectification Index: two-tailed paired t-test; EPSC ampltiudes: Wilcoxon matched-pairs signed rank test.*

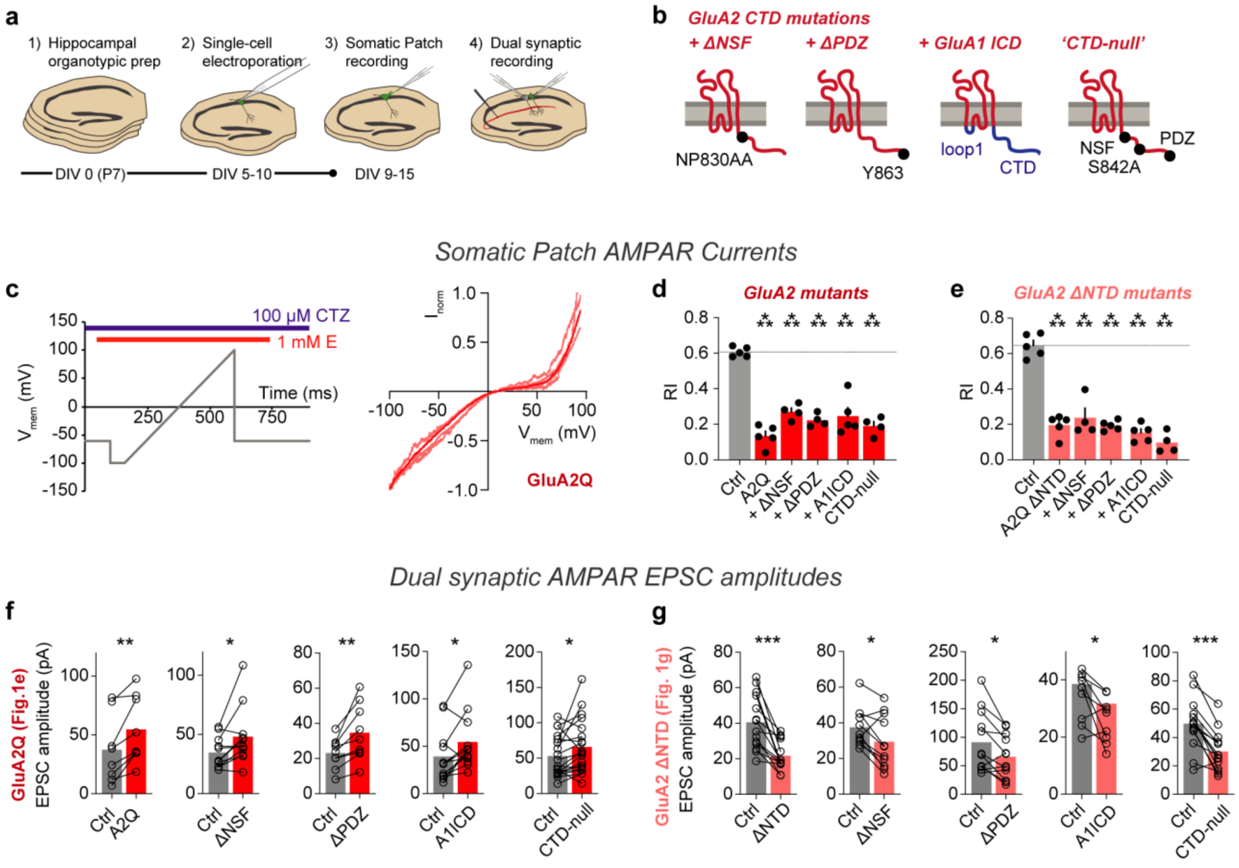


**Supplementary Figure 1. Somatic and Synaptic recordings on GluA2Q expression. a** Approach for analysis of mutant AMPAR constructs. **b** Construct schematics depicting the locations of CTD mutations on GluA2 ΔNTD. Peptide topology is displayed relative to the membrane (grey). **c** Somatic AMPAR currents demonstrate surface trafficking for GluA2 mutations by rectification index measurements. Protocol for surface rectification measurement by glutamate (E) and cyclothiazide (CTZ) application during a membrane voltage ramp (left). Representative glutamate gated currents for GluA2 overexpressing neurons (right, individual cells - light, average response - bold), normalised to -100 mV amplitude. **d** Somatic patch rectification index on overexpression of CTD mutation constructs of GluA2Q (*Ctrl:* 0.61 ± 0.01, n = 5; *GluA2Q:* 0.13 ± 0.03, n = 5; *GluA2Q ΔNSF:* 0.27 ± 0.02, n = 4; *GluA2Q ΔPDZ:* 0.22 ± 0.02, n = 4; *GluA2Q A1ICD:* 0.25 ± 0.05, n = 5; *GluA2Q CTD-null:* 0.19 ± 0.03, n = 4; [F(5, 21) = 36.38, p<0.0001]), and **e,** GluA2Q ΔNTD (*Ctrl:* 0.65 ± 0.03, n = 5; *GluA2Q ΔNTD:* 0.20 ± 0.03, n = 5; *GluA2Q ΔNTD ΔNSF:* 0.24 ± 0.05, n = 4; *GluA2Q ΔNTD ΔPDZ:* 0.19 ± 0.01, n = 5; *GluA2Q ΔNTD A1ICD:* 0.15 ± 0.02, n = 5; *GluA2Q ΔNTD CTD-null:* 0.10 ± 0.03, n = 4; [F(5, 22) = 46.22, p<0.0001]). **f-g** Dual synaptic EPSC amplitudes for normalized data presented in **Figure 1e** and **1g**. Data values are presented in **Supplementary Table 1**.

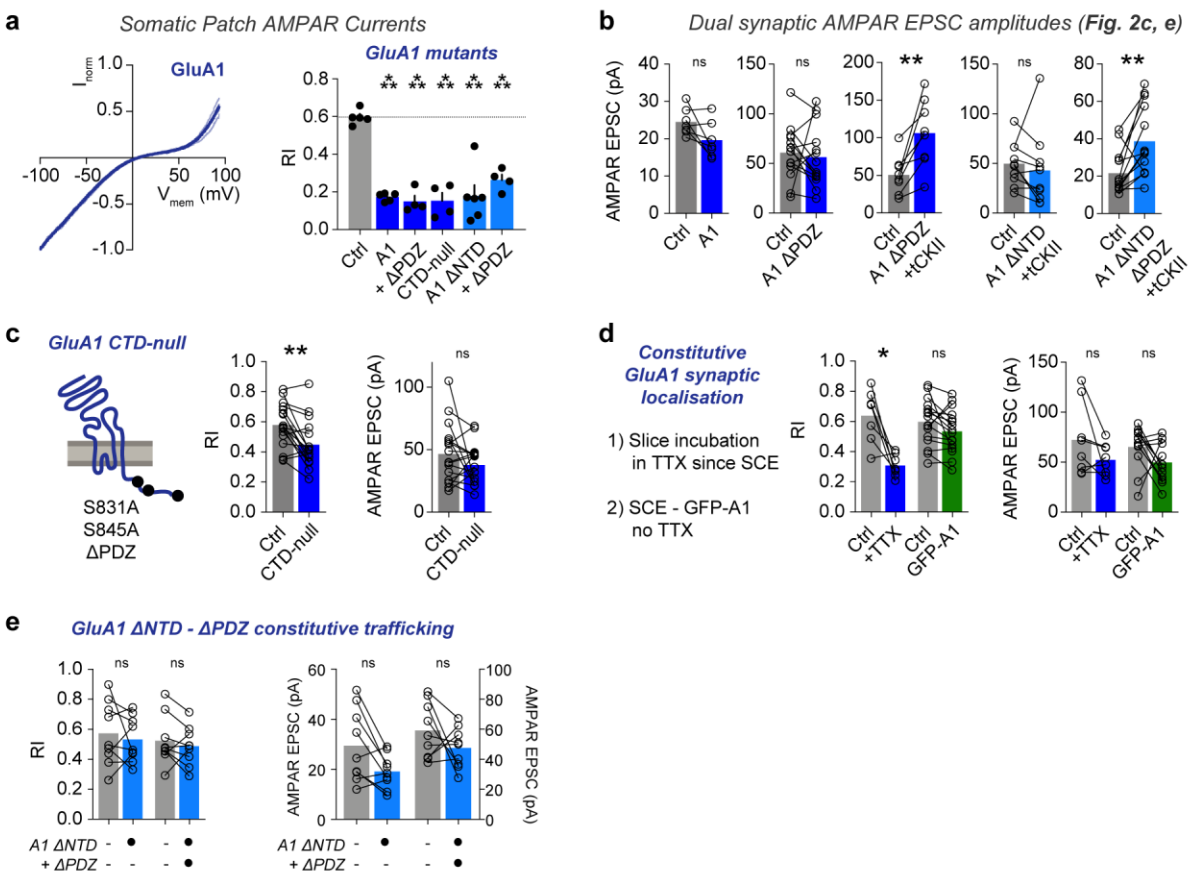


**Supplementary Figure 2. Somatic and Synaptic recordings on expression of GluA1. a** Somatic AMPAR currents demonstrate surface trafficking for GluA1 mutations by rectification index measurements GluA1 (*Ctrl:* 0.60 ± 0.02, n = 5; *GluA1:* 0.17 ± 0.01, n = 5; *GluA1 ΔPDZ:* 0.15 ± 0.03, n = 4; *GluA1 CTD-null:* 0.15 ± 0.04; *GluA1 ΔNTD:* 0.18 ± 0.06, n = 6; *GluA1 ΔNTD ΔPDZ:* 0.26 ± 0.03, n = 4; [$F(5, 22) = 21.73$, $p<0.0001$]). **b** Synaptic EPSC amplitudes for normalized data in **Figures 2c** and **2e**. **c** Schematic and synaptic recordings demonstrating synaptic localisation of GluA1 CTD-null receptors. **d** GluA1 constitutive synaptic localization does not require ongoing slice activity, as RI is altered even which activity is blocked by 1 µM TTX application from the day of transfection. Synaptic localization of GluA1 is prevented by EGFP-tagging at the N-terminus. **e** GluA1 ΔNTD does not contribute to synaptic transmission regardless of $PDZ_{GluA1}$ interactions. Data values are presented in **Supplementary Table 2**.

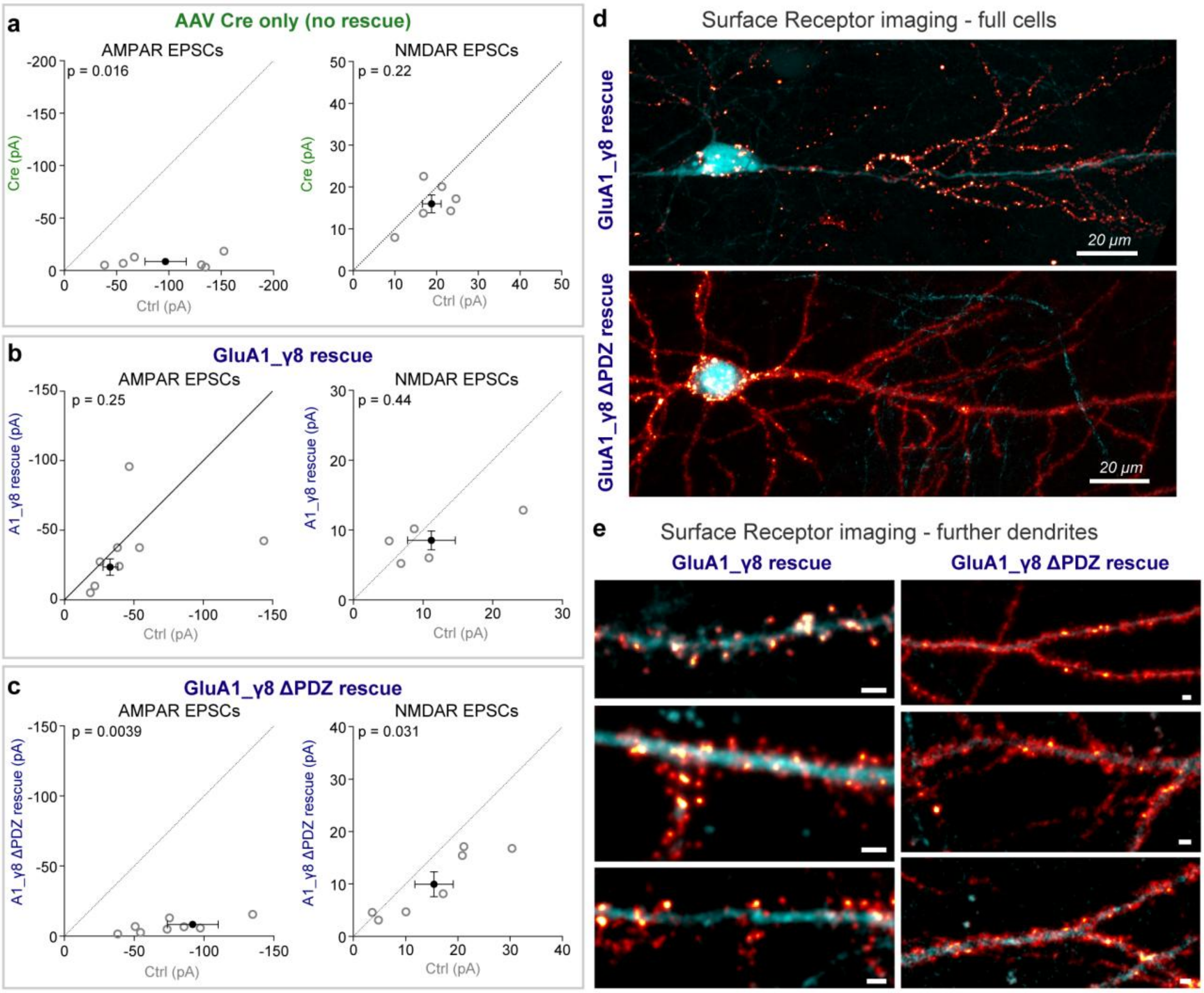


**Supplementary Figure 3. Supporting data for knockout and rescue of GluA1_ γ8 receptors. a-c** Scatter graphs of synaptic AMPAR and NMDAR EPSC amplitudes between Cre-transduced and untransduced (Ctrl) neurons (**a** - AMPAR EPSCs - *Ctrl:* -86.5 ± 19.6 pA, *Cre:* -7.9 ± 2.2 pA, n = 7 pairs, p=0.016; NMDAR EPSCs - *Ctrl:* 22.7 ± 5.4 pA, *Cre:* 18.3 ± 3.3 pA, n = 6 pairs, p=0.22), control and GluA1_γ8 rescued pairs (**b -** AMPAR EPSCs - *Ctrl:* -48.8 ± 14.3 pA, *GluA1_γ8 rescue:* -35.1 ± 9.9 pA, n = 8 pairs, p=0.25; NMDAR EPSCs - *Ctrl:* 11.2 ± 3.4 pA, *GluA1_γ8 rescue:* 8.6 ± 1.4 pA, n = 5 pairs, p=0.44), and control and GluA1_γ8 ΔPDZ rescued pairs (**c -** AMPAR EPSCs - *Ctrl:* -91.8 ± 18.2, *GluA1_γ8 ΔPDZ:* -8.5 ± 2.0, n = 9 pairs, p=0.0039; NMDAR EPSCs - *Ctrl:* 15.5 ± 3.7 pA, *GluA1_γ8 ΔPDZ:* 10.0 ± 2.4 pA, n = 7 pairs, p=0.031). **d-e** Images of surface AMPARs cell distribution on GluA1_γ8 rescue ± $PDZ_{TARP}$ interactions (cyan - tdTomato cell filler, red/glow - Streptavidin647 labelled surface AMPARs). Dendrite scale bars = 4 μm.

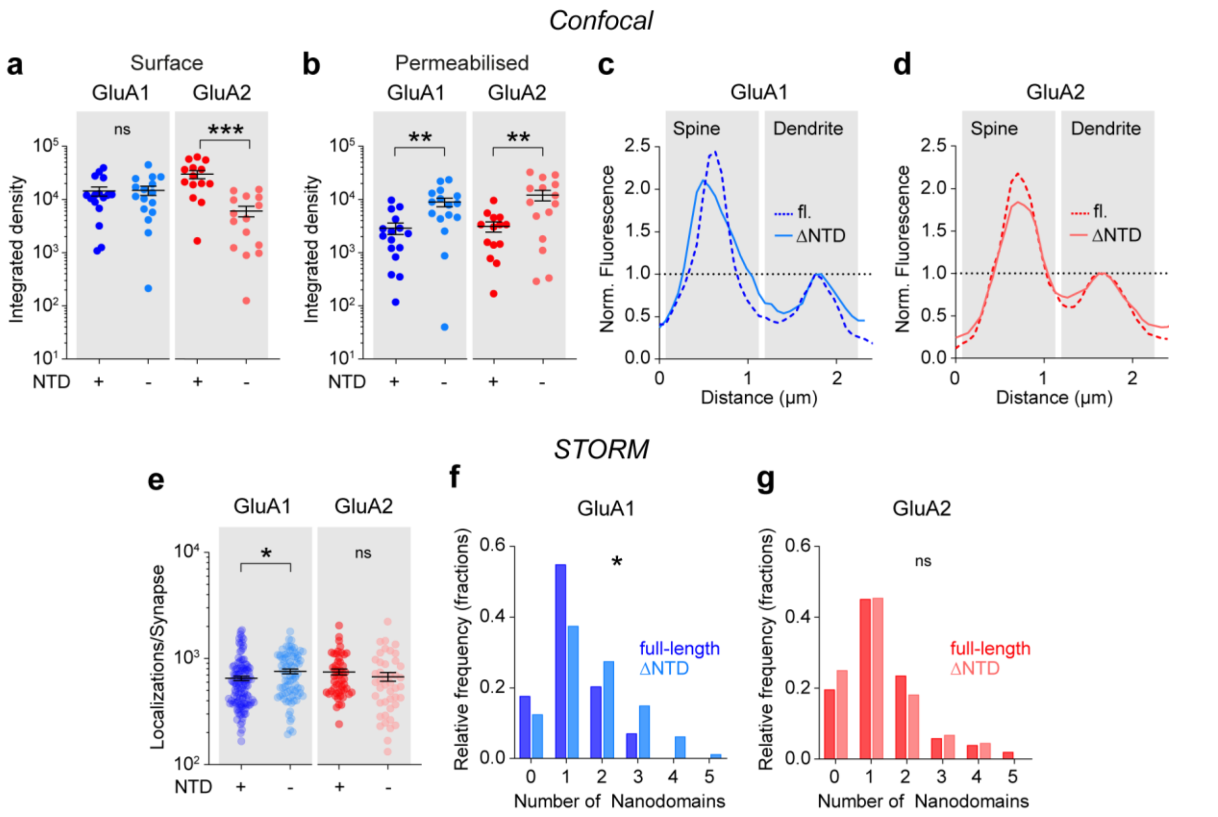


**Supplementary Figure 4. Supporting data for confocal and STORM imaging of cultured hippocampal neurons. a** Surface expression (confocal imaging) was unaltered for GluA1 ΔNTD receptors (integrated densities, left, *GluA1 full-length:* 14410 ± 2817, 16 cells, *GluA1 ΔNTD*: 14880 ± 2891; 16 cells, 5 independent preparations; p=0.9080, unpaired t-test) but significantly reduced for GluA2 ΔNTD receptors (integrated densities, right *GluA2 full-length:* 29855 ± 5081, 14 cells, *GluA2 ΔNTD*: 6072 ± 1360; 15 cells, 4 independent preparations; p < 0.0001, unpaired t-test) compared to respective full-length controls. **b** NTD deletion resulted in increased intracellular expression levels for both GluA1 (integrated densities, left, *GluA1 full-length:* 4007 ± 956.1, *GluA1 ΔNTD*: 12403 ± 2352; n-numbers given in a; p=0.0025, unpaired t-test) and GluA2 (right, *GluA2 full-length:* 4295 ± 923.4, *GluA2 ΔNTD*: 16793 ± 3806; n-numbers given in a; p=0.0046, unpaired t-test) compared to full-length receptors. **c-d** Representative line-scans across spines and dendrites for full-length and NTD-deleted GluA1 (**c**) and GluA2 (**d**) receptors (See Figure 4). **e** NTD deleted GluA1 (left, *GluA1 full-length: 647.5* ± 33.8, *GluA1 ΔNTD* 760.6 ± 39.5; p=0.0190, unpaired t-test) but not ΔNTD GluA2 receptors (right, *GluA2 full-length: 749.9* ± 50.4, *GluA2 ΔNTD:* 675.4 ± 64.9; p=0.0838, unpaired t-test) displayed more localisations per synapse upon 3D STORM imaging. **f-g** Relative frequency distribution of nanocluster numbers across individual synapses was changed in GluA1 ΔNTD receptors (left, p = 0.017, Kolmogorov-Smirnov test) but unaffected in NTD-lacking GluA2 receptors (right, p>0.999,

Kolmogorov-Smirnov test). Data in panels a, b, and e shown as mean ± SEM. n-numbers for STORM data are given in legend to Figure 5.

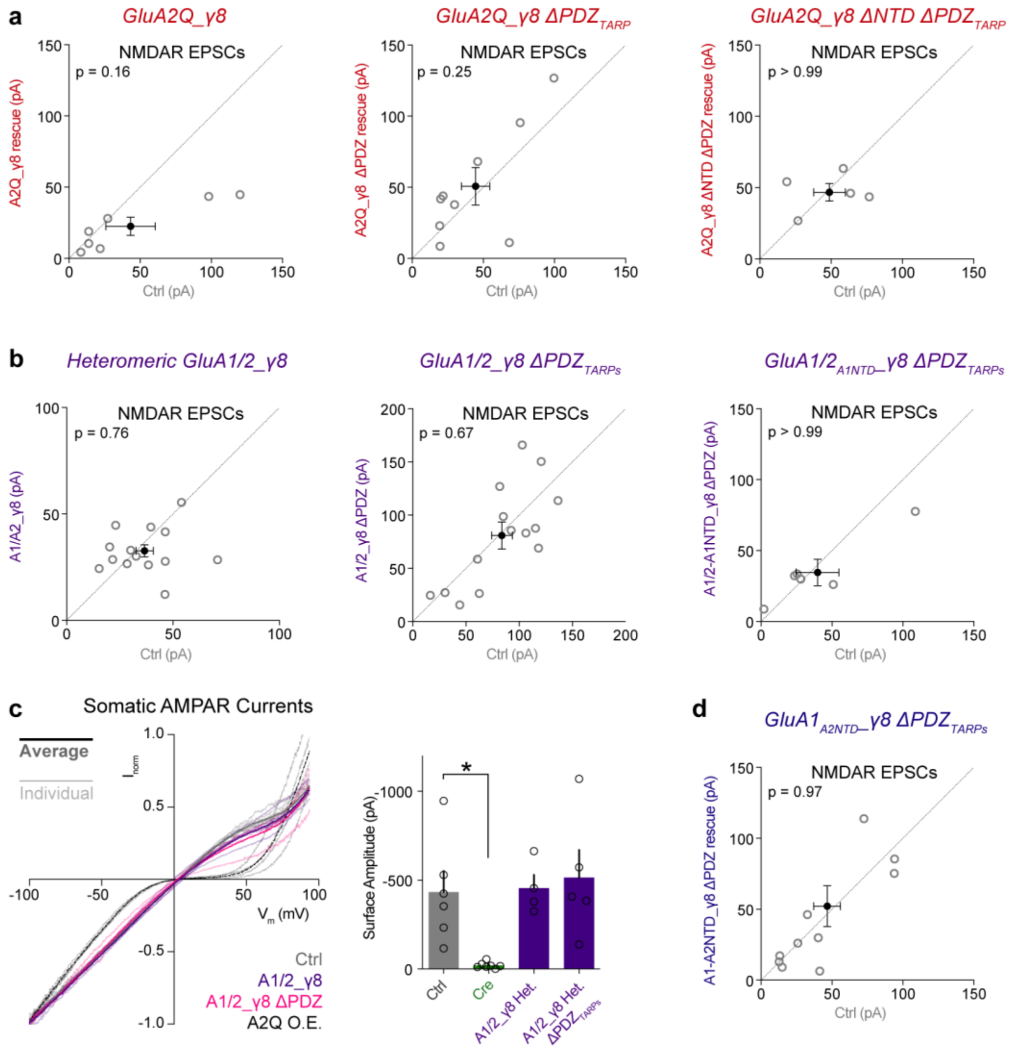


**Supplementary Figure 5. Supporting figures for knockout and rescue using GluA2_γ8 or heteromeric AMPARs. a** Synaptic NMDAR currents are unchanged after AMPAR knockout and rescue with different GluA2Q_γ8 constructs (*GluA2Q_γ8 - Ctrl:* 43.4 ± 17.3 pA, *Rescue:* 22.5 ± 6.4 pA, n = 7 pairs, p=0.16; *GluA2Q_γ8 ΔPDZ - Ctrl:* 44.6 ± 10.0 pA, *Rescue:* 50.8 ± 13.2 pA, n = 9 pairs, p=0.25; *GluA2Q_γ8 ΔPDZ ΔNTD - Ctrl:* 48.9 ± 11.1 pA, *Rescue:* 46.8 ± 6.1 pA, n = 5 pairs, p>0.99). **b** Synaptic NMDAR currents are unchanged after AMPAR knockout and rescue with different tandem heteromeric receptors (*A1_γ8/A2_γ8 - Ctrl:* 36.7 ± 4.1 pA, *Rescue:* 32.7 ± 2.9 pA, n = 14 pairs, p=0.76; *A1_γ8/A2_γ8 ΔPDZs - Ctrl:* 83.9 ± 9.7 pA, *Rescue:* 81.0 ± 12.8 pA, n = 14 pairs, p=0.67; *A1_γ8/A2$_{A1NTD}$_γ8 ΔPDZs - Ctrl:* 39.8 ± 15.2 pA, *rescue:* 34.7 ± 9.3 pA, n = 6 pairs, p>0.99). **c** Somatic AMPAR currents from outside-out patches on heteromeric receptor rescue. *left* - Normalised current traces from holding potential ramp application to patches in the presence of glutamate, demonstrating non-rectifying responses on heteromeric receptor rescue. Both cell-averaged responses (bold) and individual cells (transparent) are shown (untransfected cells - grey,

A1_γ8/A2_γ8 rescue - purple, A1_γ8/A2_γ8 ΔPDZs rescue - pink) against strongly rectifying GluA2Q overexpressing cell responses (black) for reference. *right* **-** Current amplitudes from somatic patches demonstrate abolishment of AMPAR response on Cre transduction, and similar surface receptor rescue by heteromeric receptors with and without TARP PDZ interactions (*Ctrl:* -434 ± 118 pA, n = 6; *Cre:* -20 ± 6.5 pA, n = 7; *A1_γ8/A2_γ8 rescue:* -456 ± 74 pA, n = 4; *A1_γ8/A2_γ8 ΔPDZs rescue:* -515 ± 155 pA, n = 5; one-way ANOVA with Dunnett' multiple comparisons test [F(3,18) = 6.007, p=0.0051]). **d** NMDAR currents are unchanged on rescue with $GluA1_{A2NTD}$_γ8 ΔPDZs (*Ctrl:* 46*.7* ± 9.4 pA, *Rescue:* 52.3 ± 14.5, n=11 pairs, p=0.96).
